# Programmable DNA Protonuclei Reveal Environmental Context on Protein Phase Separation

**DOI:** 10.64898/2026.04.07.716875

**Authors:** Johann Fritzen, Avik Samanta, Nele S. Kuhr, Antonia Preuß, Erin L. Sternburg, Lukas S. Stelzl, Jasper J. Michels, Dorothee Dormann, Andreas Walther

## Abstract

Understanding protein phase separation in cellular environments remains a major challenge, as *ex vivo* assays often fail to capture the influence of environmental context – such as crowding, multimodal interactions, and the dynamic properties of the cytosol or nucleus. Here, we introduce programmable DNA-based protonuclei (PN) as nucleus-inspired compartments to probe phase separation of the neurodegeneration-linked protein FUS. We show that FUS partitioning and condensate formation are highly sensitive to nucleic acid sequence, spatial confinement, and viscoelastic properties of the PN core. Notably, classical test-tube affinity assays fail to predict protein behavior within the crowded and multivalent PN environment. By tuning DNA crosslinking, we modulate condensate dynamics and suppress liquid-to-solid transitions of FUS – a hallmark of disease. These findings demonstrate that multivalent, confined environments fundamentally reshape protein-nucleic acid interactions and phase behavior. The PN platform complements test-tube assays and complex cellular settings and enables to dissect nuclear condensates under controllable conditions.

## Introduction

Phase separation (PS) leading to biomolecular condensates plays a key role in cell regulation, organization, and stress response.^1–3^ Many of such condensates are formed by protein and nucleic acid components, for instance, the nucleolus, Cajal bodies, paraspeckles, and nuclear speckles in the cell nucleus, and stress granules or P-bodies in the cytoplasm. ^4^ Understanding such protein/nucleic acid condensates is of critical importance to further our knowledge about disease development and cellular regulatory processes, which is particularly challenging for condensates formed by multiple components and occurring in complex and nucleic acid-rich environments, such as the nucleus. Indeed, the nucleus presents a unique environment comprising numerous RNA/protein- and chromatin-based condensates whose assembly and material properties are governed by high nucleic-acid content, transcriptional activity, and viscoelastic chromatin constraints.^4,5^

One key problem is the translatability of facile test-tube solution experiments to *in cellulo* situations.^6^ Simple solution investigations benefit from high throughput and widely tunable conditions, and have the potential to speed up mechanistic understanding due to reductionist settings. However, such systems rarely capture the complexity of the cellular environment, which is not only crowded but also compositionally and mechanically heterogeneous. This discrepancy limits the translatability of insights gained from dilute solution assays to real biological function. The addition of molecular crowders has been one approach to arrive at situations mimicking more closely the crowded cellular environment. Nonetheless, some crowders influence PS beyond their crowding contribution,^7,8^ partition into condensates^9–11^ or change condensate properties at high concentrations.^12^ Beyond these limitations, such conditions cannot replicate anything vaguely reminiscent of the cell nucleus, for which it would be very beneficial to have tunable *ex cellulo* model systems.

A particular gap exists in understanding how nucleic acid sequence, spatial confinement, and material state jointly shape PS of biopolymers – a triad that remains unaddressable thus far in conventional approaches. A system that allows precise control over these factors could provide critical insights into the mechanisms of nuclear PS, especially for proteins that are implicated in disease, such as neurodegenerative diseases and cancer, through misregulated condensation.^13,14^ Importantly, several RNA-binding proteins implicated in neurodegenerative diseases, such as amyotrophic lateral sclerosis (ALS) and frontotemporal dementia (FTD), including Fused in sarcoma (FUS) and TDP-43, form nuclear condensates *in vivo*^15^ and occasionally form pathological aggregates in the nucleus.^16^ FUS undergoes PS at sites of DNA double-strand breaks, where FUS-dependent PS is required for the recruitment of DNA repair factors and higher-order clustering of chromatin nanodomains. This identifies chromatin-associated, nucleic acid-rich and mechanically constrained nuclear environments as critical regulators of FUS condensate function.^17^

We hypothesized that a critical cross-fertilization of the field of biological PS would be possible by transferring concepts from all-DNA artificial cells^18–22^ to study condensate formation processes in tailored nucleus-inspired environments. Such synthetic structures are built entirely from DNA and are tunable in composition, structure and viscoelastic properties. To this end, we introduce here a protonucleus (PN) platform as an advanced and versatile test system between classical test tube/solution experiments and *in cellulo* studies. We explore this system for a model protein implicated in neurodegenerative disease – the FUS protein – which is known to undergo PS and to bind to nucleic acids.^23–26^ We find that the recruitment and PS of FUS is not only highly dependent on DNA and RNA sequence, but that classical *ex vivo* methods such as electrophoretic mobility shift assays (EMSA) cannot fully predict the interactions found in crowded and restrained DNA PN environments. Additionally, we show that the dynamic/viscoelastic properties of the PN significantly influence PS of FUS to the point of full suppression. Our new *in protonucleo* method based on a tunable DNA nanoscience approach introduces a highly versatile platform that complements simple in solution *ex vivo* assays and *in cellulo* studies.

## Results and Discussion

### Synthetic all-DNA PN as a protein PS platform

The PN builds on our work on DNA-based core-shell particles formed by temperature-induced PS of two long ssDNA multiblock copolymers, namely p(A_20_-m) and p(T_20_-n).^19,21,22,27–29^ Therein, A_20_ and T_20_ correspond to homo-repeats of adenine and thymine nucleobases, respectively, and m and n are ssDNA barcodes that allow for functionalization (sequences in Scheme 1a, Supplementary Table 1). In short, PN form by selective PS of p(A_20_-m) during heating, whereafter p(T_20_-n) hybridizes during cooling onto the phase-separated droplets, stabilizing them at the core/shell interface by multivalent A_20_/T_20_ duplexes (Scheme 1a). The resulting PN has a kinetically trapped, liquid pool of long ssDNA inside. The DNA concentration is 5-13 g/L, which approximates that of the nucleus in a cell of about 9-31 g/L (Supplementary Table 2). The PN shell is highly permeable and allows for the uptake of macromolecules, such as proteins.^27^

To apply the PN system to a biologically and disease-relevant model system, we selected the FUS protein because it is a very well-studied nucleic acid-binding protein.^25,26,30^ In cells, FUS is involved in splicing, DNA damage repair, transcription regulation, RNA transport and translation regulation.^31–34^ FUS is mostly located in the nucleus of healthy cells, but can be mislocalized to the cytoplasm of nerve cells because of mutations in the nuclear localization signal (NLS), or other defects to Transportin-1 (TNPO1)-mediated nuclear import of FUS (Scheme 1b).^33,35–37^ Cytosolic FUS phase separates over time and FUS condensates undergo an irreversible liquid-to-solid transition to a non-functional amyloid-like state,^23,38–40^ implicated in neurodegenerative diseases, such as ALS and FTD.^41–43^

**Scheme 1:**
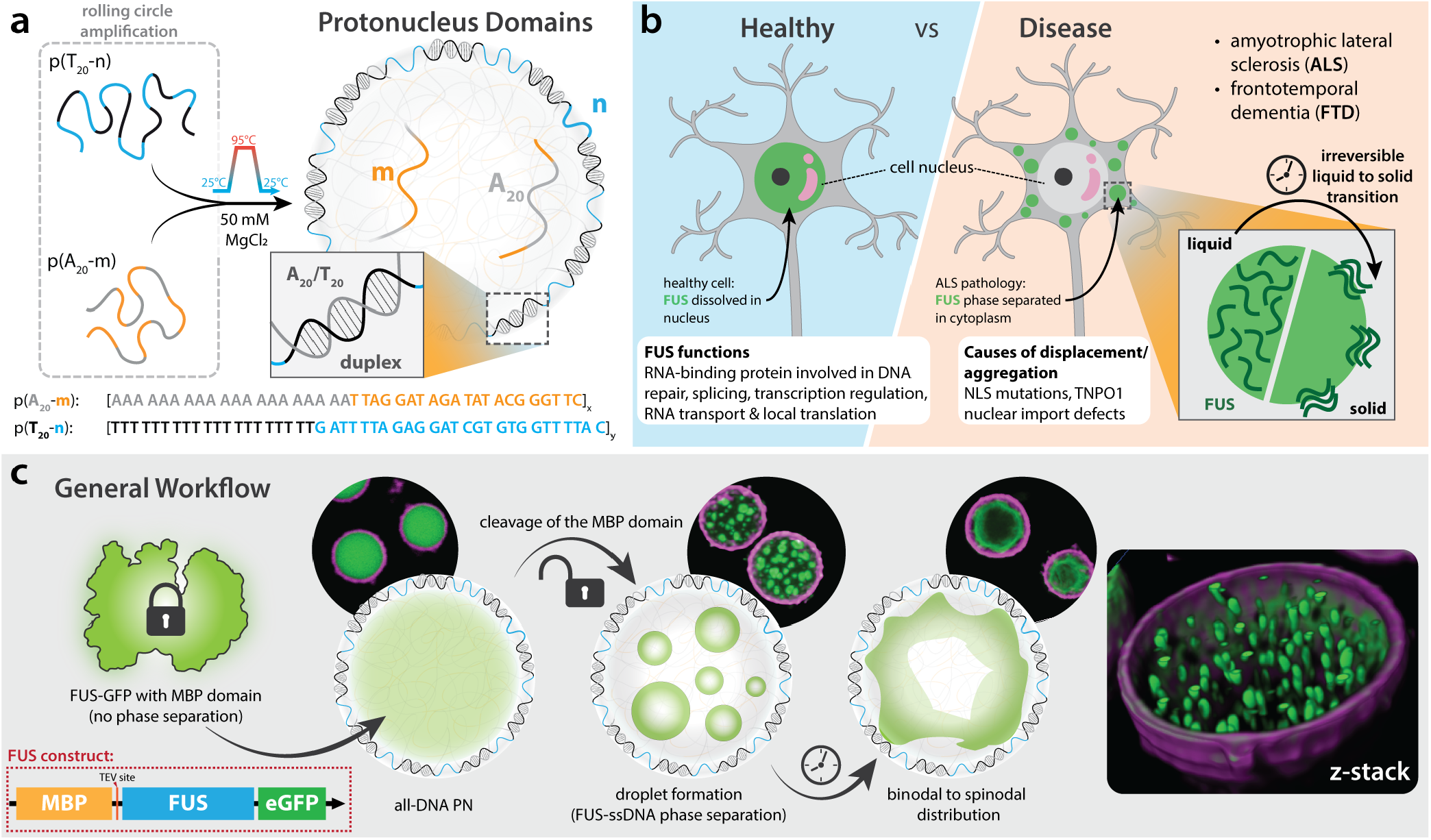
PN platform and biological relevance of FUS PS. **a** Simplified preparation of our PN platform and the relevant addressable domains m, n and A_20_. **b** Schematic comparison of FUS in a healthy neuron to the FUS pathology in ALS and FTD. **c** The general workflow of a PS experiment with MBP-FUS-GFP (soluble) and all-DNA PN, encompassing protein loading, MBP-tag cleavage, and microscopic visualization of FUS PS inside PN.

Scheme 1c visualizes the general workflow. We use recombinantly produced full-length FUS with a C-terminal eGFP tag for fluorescence imaging by confocal laser scanning microscopy (CLSM), as well as an N-terminal protease-cleavable maltose binding protein (MBP) tag for enhanced solubility and initial suppression of PS. This strategy allows to introduce MBP-FUS-GFP into PN in a soluble state, and, once the sample is equilibrated, to trigger PS of FUS by cleaving off the MBP tag with TEV protease. Control experiments using isolated MBP and TEV protease confirm that these components and their combination do not induce PS of the PN core (see Supplementary Fig. 1-3).

### FUS loading ratio controls PS morphology

We first investigate the influence of FUS-GFP concentration on the PS behavior. In these experiments we use pristine PN only labeled at their shell with n*^Atto565^ to visualize the shell in CLSM (magenta channel). The ratio of FUS-GFP to repeating units (r.u.) of p(A_20_-m) describes the increased loading of FUS-GFP into the PN. Partitioning values for each loading ratio (LR) and fluorescence recovery after photobleaching (FRAP) of FUS-GFP before MBP-cleavage are shown in SI Section 4 (Supplementary Fig. 4 and 5). Figure 1 depicts three major domain windows that occur at different FUS-GFP/p(A_20_-m) ratios of 0.10 to 1.50. We have chosen representative snapshots of intermediate morphologies (0-30 min) and end state morphology (5 h) of the FUS PS process. At a low LR (≲0.30), phase-separated FUS-GFP droplets appear within the PN, resembling a binodal domain. Binodal PS takes place in the metastable boundary region of the phase diagram. It requires fluctuations to nucleate PS, which usually leads to a droplet morphology by growth and coalescence. In contrast, spinodal decomposition happens spontaneously and rapidly for more concentrated solutions in the center of the phase diagram. This typically leads to co-continuous morphologies that coarsen and can also form a single domain at the end of the PS process.^44–47^ Interestingly, for LRs between approximately 0.60 and 0.90, a co-continuous morphology occurs that is reminiscent of the spinodal domain. Key morphological features include surface-directed spinodal decomposition, characterized by formation of a wetting layer at the p(A_20_-m)/p(T_20_-n) interface followed by a depletion layer. This pattern repeats throughout the condensate, generating co-continuous structures with a characteristic length scale. Over time, continued coarsening and merging of these domains give rise to a single condensate (Figure 1, end state). Finally, at high LRs (≳ 1.20) the dilute phase forms droplets within the dense phase, marking the inverted region of the phase diagram.

**Figure 1:**
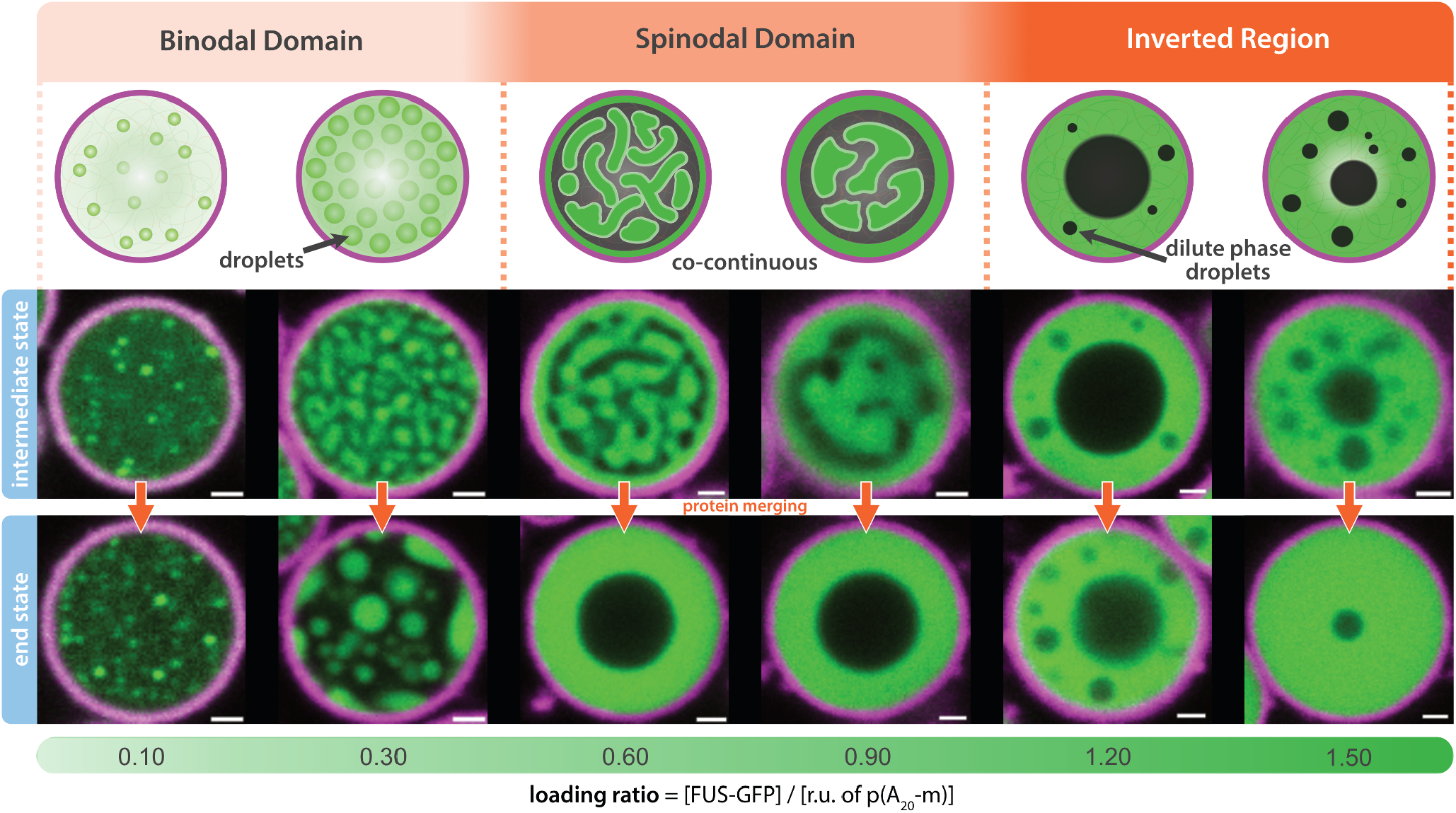
Different ratios of FUS-GFP to core repeating unit of p(A_20_-m) of PN lead to diverse PS morphologies. The PN shell is shown in the magenta channel, FUS-GFP is in green. Imaging of the samples was done continuously for the initial 30 min, with representative snapshots shown as “intermediate state”. Snapshots shown as “end state” are at 5 h after TEV addition. Overview images are shown in Supplementary Fig. 6. Additionally, the initial PS can be viewed in Supplementary Movie 1. The repeat sequence of p(A_20_-m) is shown in Scheme 1a. Scale bar = 1 µm.

### Continuum model predicts FUS phase behavior in PN

To rationalize and predict FUS phase behavior in multicomponent mixtures, we next introduce a continuum model. The model (details in Section 5 of the SI) treats the PN and their contents as order parameter fields, whose spatiotemporal variations are governed by minimization of a free energy. For a solution containing *m* - 1 solutes (*m* indexing the solvent) in the presence of *n* PN, this free energy is written as a function of sets of conserved (volume fractions) {*c*_1 ≤*i*≤*m*_ (*r*)} and non-conserved order parameters {*φ*_1 ≤*k* ≤*n*_ (*r*)} as:

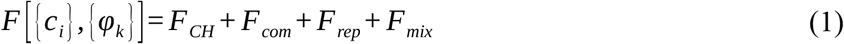

The order parameter *φ*_*k*_ represents the *k*^th^ PN and assumes values of 1, 0 and 0 < *φ*_*k*_ < 1 in its interior, exterior and boundary region. Equation (1) comprises a Cahn-Hilliard free energy (*F*_*CH*_), a contribution due to PN compressibility (*F*_*com*_), a penalty (*F*_*rep*_) associated with spatial overlap between PN, and a Flory-Huggins-type mixing free energy (*F*_*mix*_), which are detailed in Section S5.1 of the SI. The evolution equations for the concentrations and PN are:

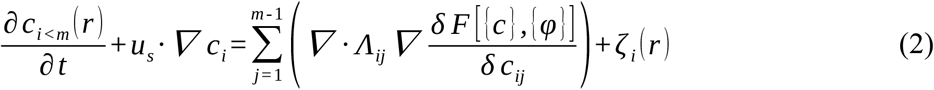

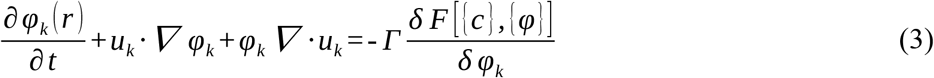

with *u*_*k*_ (*r*) the local velocity in each phase field and *u*_*s*_ the divergence-free (solenoidal) component of the total velocity field 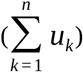. Under the assumption of overdamped dynamics, we relate the local velocity to a thermodynamic restoring force as 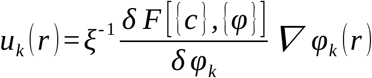, with *ξ* a friction coefficient associated with PN motion. *Γ* and *Λ*_*ij*_ are kinetic and mobility coefficients, governing the dynamics of, respectively, the non-conserved and conserved order parameter fields, the latter being subject to delta correlated noise represented by *ζ* _*i*_ (*r*), numerically implemented as described previously^48,49^.

By numerically solving the dynamic Equations (2) and (3), we simulate FUS co-PS in PN at various LRs. A calculation comprises two stages: (i) a generation stage and (ii) a quenching stage. In the generation stage, the MBP-FUS loaded PN are formed. To this end, we nucleate the PN in a homogeneous ternary mixture comprising generic “DNA” (e.g. p(A_20_-m)), “MBP-FUS” and “solvent”, whereupon the components partition across the PN exterior (α-phase) and interior (β-phase), dictated by binary interaction parameters 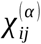 and 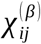. DNA strongly partitions inside the PN as we set 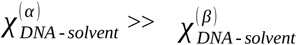. In contrast, the MBP-FUS is expected to have the same interaction with the solvent inside and outside the PN, so 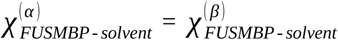 but should preferentially locate inside the PN on account of its DNA binding property: 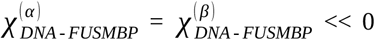. Uptake of MBP-FUS is further driven by assuming marginal solvent conditions, evaluating 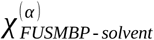 and 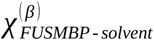 just below their critical value. All values of the interaction parameters in the generation stage are given in Figure 2a and listed in Supplementary Tables 3 and 4.

**Figure 2:**
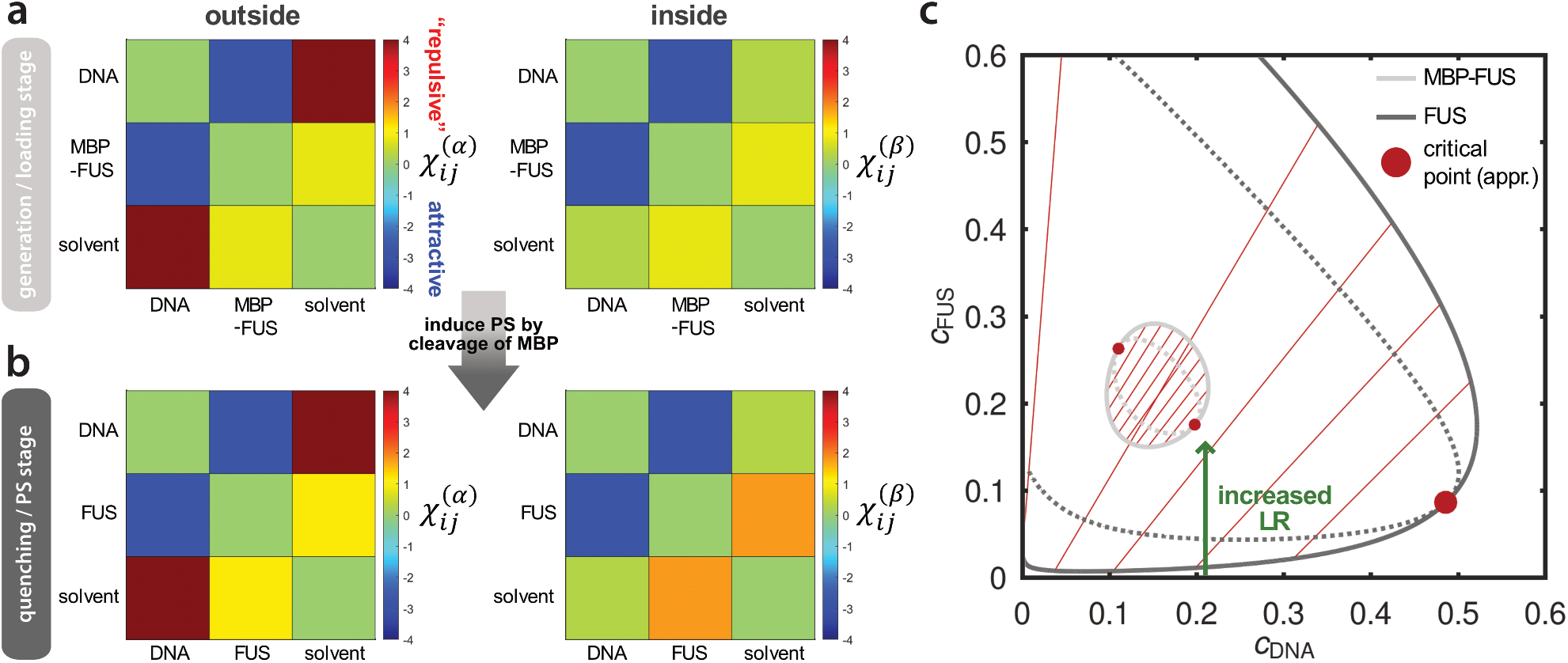
Interaction matrices and calculated ternary phase diagrams. **a** and **b** color scale representation of the Flory-Huggins interaction parameters in the generation stage and quenching stage, respectively, i.e., for DNA:MBP-FUS:solvent and DNA:FUS:solvent outside (left) and inside (right) the PN. **c** Ternary phase diagrams, calculated by equating chemical potentials and osmotic pressures in/of coexisting phases using previously reported methods.^50–52^ The diagrams show the considerable increase of the miscibility gap upon cleaving the MBP tag: the light gray and dark gray phase diagrams respectively correspond to DNA:MBP-FUS:solvent and DNA:FUS:solvent (spinodal: dotted lines, binodal: solid lines). The solid green arrow indicates the approximate core concentration in each PN (0.21) and how the LR increases with the FUS concentration inside the PN.

In the quenching stage we numerically simulate the co-PS of FUS and DNA upon removing the MBP tag. We directly target correct co-PS, because, as we will see below, DNA in fact co-phase separates with FUS in the PS process in pristine PN. MBP removal is achieved by subjecting the loaded PN formed in the generation stage to a stepwise increase in *χ* _*FUSMBP* - *solvent*_, which logically becomes *χ* _*FUS* - *solvent*_ (see Figure 2b and Supplementary Table 3). For simplicity, we do not consider the cleaved MBP tag as an additional component, nor do we consider changes in molecular volume. Naturally, the increase applies to the interior as well as the exterior of the PN and causes a drastic change in the phase diagram: the small miscibility gap encountered for DNA:MBP-FUS:solvent widens significantly (see Figure 2c), pushing the phase boundaries in opposite directions. This gives rise to coexistence between a dense phase, enriched in FUS + DNA and a solvent-rich phase nearly depleted in FUS but containing a fraction of DNA. As a result of the expansion in the miscibility gap, compositions that were formerly in a single-phase region now find themselves in a coexistence region, which leads to PS inside the PN.

Figure 3a and Supplementary Movie 2 show how three PN nucleated in a homogeneous solution of DNA and MBP-FUS grow to reach their equilibrium size while repelling each other to avoid spatial overlap. Figure 3b shows numerically simulated phase-separated early-time morphologies in the quenching stage for a constant mean DNA concentration of 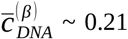 (see green arrow in Figure 2c) and increasing loading of FUS(-MBP) (left to right). The evolution of the morphologies is in agreement with the experiments (Figure 1). When increasing the LR, the dense phase first organizes in droplets, then becomes a co-continuous regime, before crossing over into a phase-inverted morphology for LR >> 1. This corresponds to the binodal, spinodal and inverted phase regimes. Furthermore, equally in agreement with the experiments, assuming a mild short-range attraction between DNA and the interior wall of the PN (details in Section 5.2 of the SI), gives rise to a surface enrichment of the dense phase and surface directed PS (see e.g., LR = 0.8, 1). Collectively, these numerical simulations explain and fundamentally validate the experimentally observed morphologies on account of solid PS theory in multicomponent mixtures.

**Figure 3:**
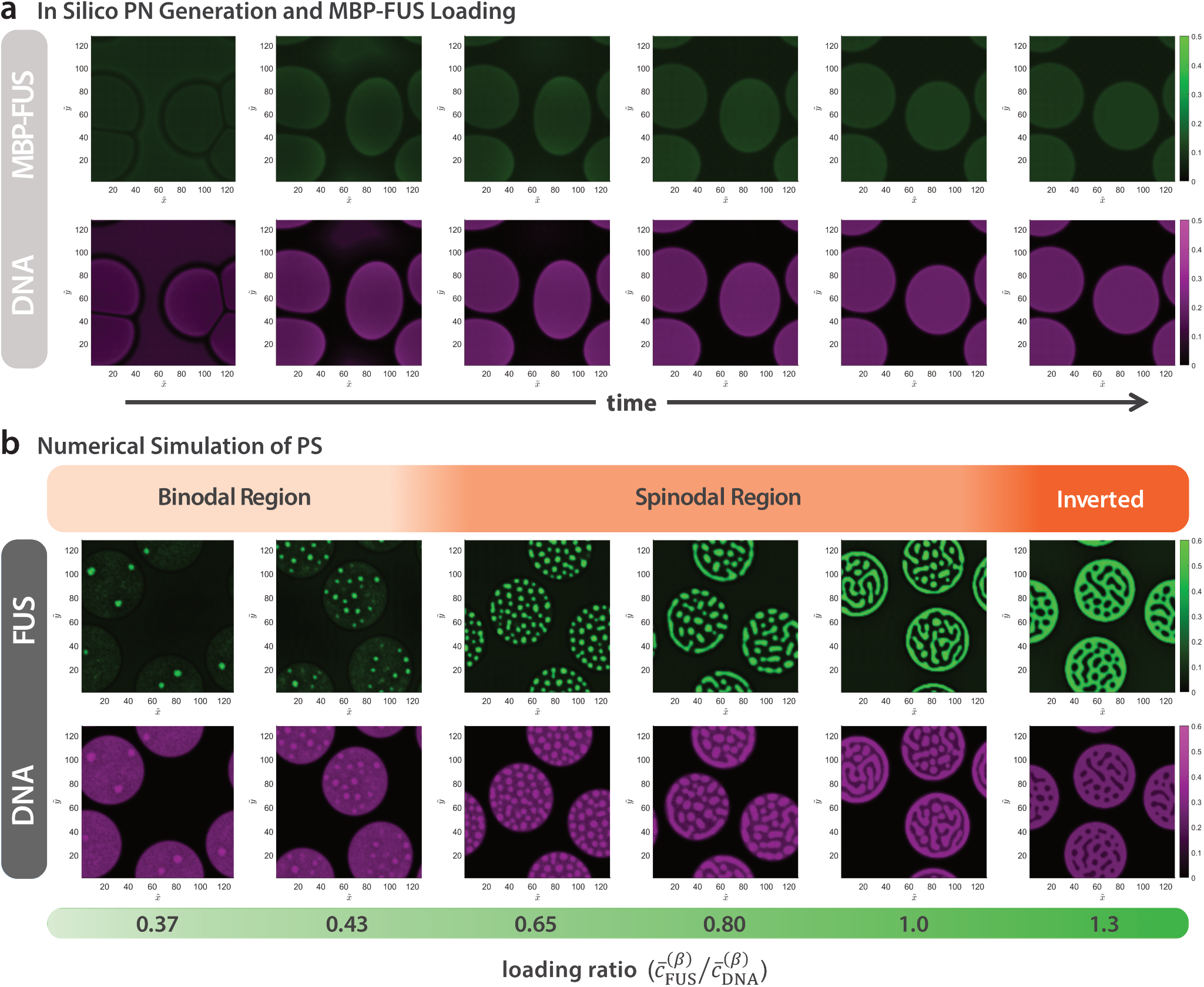
Numerical simulations. **a** Exemplary PN generation at LR = 0.65 (see Supplementary Movie 2). Droplets become more spherical due to ongoing PS and partitioning of MBP-FUS. The time domain is 2048 dimensionless time units. **b** PS end stages with final morphology snapshot of DNA:FUS co-PS inside PN, triggered by removing the MBP tag as a function of the LR. The time domain is 40 dimensionless units for LR = 0.37 and 16 dimensionless units for the higher LRs (see Supplementary Movie 3).

### PN core sequence modifications affect FUS partitioning

One of the strengths of the PN system is the ease of functionalizing them with additional nucleic acids (either DNA or RNA oligonucleotides) via their m barcodes or A_20_ domains to understand how variations in PN core composition impact affinity and PS. This is schematically shown in Figure 4a for six samples: The pristine PN (1) is the only sample without core modification, meaning both m and A_20_ domains are not hybridized. This is different for the next two samples, where a sequence complementary to m is added (sequences in Figure 4b): (2) m*^Atto647N^ forms m/m*^Atto647N^ DNA duplexes. (3) m*-RNA^Cy5^ is a complementary RNA leading to an m/m* DNA/RNA duplex with a dangling RNA strand (39 nt in length). The remaining samples are structured analogously, with the difference of utilizing the A_20_ domain for hybridization: (4) 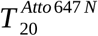 forms a simple 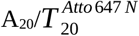 DNA duplex. (5) U_20_-RNA^Cy5^ forms a A_20_/U_20_ duplex with a dangling RNA strand. (6) T_20_-m^Atto647N^ has a dangling m as additional ssDNA overhang, effectively doubling [m] in the PN core. As RNA overhang in both RNA-containing samples, we chose a physiologically relevant RNA with high affinity to FUS.^25,53^ All these modifiers are tightly bonded to the PN via dsDNA or DNA/RNA bonding, with melting temperatures, *T*_m_,, in experimental buffer exceeding 60 °C (calculated *T*_m_ in Supplementary Fig. 7). All experiments use a LR of 0.30, hence a regime where the pristine PN/FUS-GFP system is in the binodal domain (Figure 1). We chose this concentration regime to ensure an excess of the introduced nucleic acid sequences compared to FUS-GFP.

**Figure 4:**
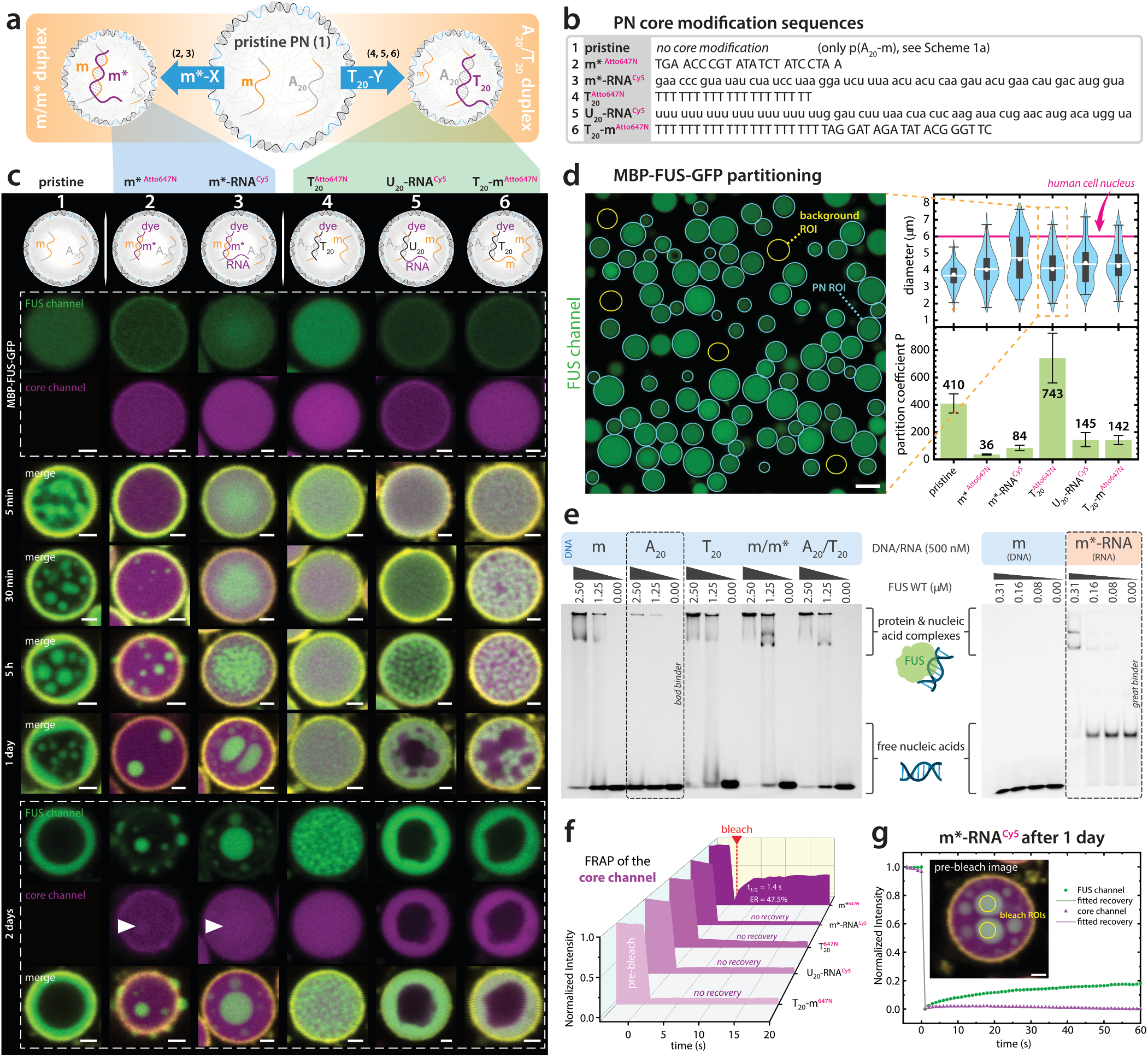
Modification of the PN core sequences alters partitioning and PS behavior. **a** Schematic illustration of the two PN core modification strategies used in the experiments. **b** DNA (2, 4, 6, uppercase) and RNA (3, 5, lowercase) sequences used for core modification. **c** CLSM Snapshots of single PN with different DNA/RNA modifications after defined time intervals of 5 min, 30 min, 5 hours, 1 day and 2 days. FUS-GFP in green, core modifications in magenta and shell in yellow channel. White arrows indicate condensates with core segregation, see Supplementary Fig. 9, 10. PN have average diameters from 3.7-4.7 µm. We scaled representative PN examples to full panel size for clarity. For overview images with indication of the selected PN, see Supplementary Fig. 11, 12. Scale bar = 1 µm. **d** Example of manual PN segmentation in sample 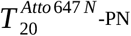 to measure PN diameter and partition coefficient P of MBP-FUS-GFP. Scale bar = 5 µm. Violin plots: diameter distribution from 2D CLSM: Box shows inter quartile range (IQR), whiskers to 1.5 x IQR, outliers in orange, mean as white line, median as white circle. Bar chart: Partition coefficient of FUS: Error bars show standard deviation. N = 146, 92, 92, 91, 83, 79 (left to right). N denotes the number of individual PN compartments. **e** EMSAs of all relevant DNA/RNA sequences to probe their binding to FUS. **f** FRAP of the pA-based core material of the PN of all samples with dye-containing core modifications before MBP-FUS-GFP addition. Note that the pristine PN (1) does not have a dye modification. See Supplementary Fig. 13, 14 for FRAP details. **g** FRAP of m*RNA^Cy5^-PN after 1 day for both the core and FUS channel. Scale bar = 1 µm. Loading ratio is 0.30 for all experiments.

Experimentally, we monitored the evolution of FUS PS over time through CLSM. Figure 4c shows single PN before TEV addition (boxed region) and at different time points after TEV addition (overview images in Supplementary Fig. 8). The mean diameters of all PN range from 3.7 to 4.7 µm (Figure 4d), thus slightly smaller but comparable in scale to human cell nuclei with an average diameter of about 6 µm.^54^

We first quantify the partitioning of MBP-FUS-GFP into PN before initiation of PS by comparing the mean GFP intensity inside the PN core with that of the solution (background; Figure 4d). The equilibrated PN show different levels of protein partitioning. Pristine PN (1) and 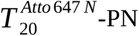 (4) exceed the other samples by far. Both RNA-containing PN (3: m*-RNA^Cy5^, 5: U_20_-RNA^Cy5^) and T_20_-m^Atto647N^-PN (6) show moderate MBP-FUS-GFP partitioning, whereas the weakest partitioning is observed for m*^Atto647N^-PN (2).

Since partitioning is expected to scale with affinity, we semi-quantitatively assessed binding of FUS to the different nucleic acid sequences using electrophoretic mobility shift assays (EMSA, Figure 4e). EMSA demonstrates a moderate affinity of FUS for all DNA sequences except A_20_, which is a poor FUS binder (Figure 4e, left). Poor A_20_-FUS binding can be explained by the strong base stacking behavior known for p(A),^55,56^ limiting its availability for FUS binding, which we also identify in atomistic molecular dynamics simulations between A_20_ and a nucleic acid-binding part of FUS (Section 11 of the SI, Supplementary Fig. 15, 16 and Supplementary Table 5). A comparison of a moderate binder (m) and a strong binder (3: m*-RNA^Cy5^) at lower FUS concentration shows the far superior binding affinity of the latter, in line with the known high RNA/FUS affinity for this particular RNA^25,53^ (Figure 4e, right).

Comparing the EMSA data to the partitioning data, some trends carry over, such as binding of FUS-GFP is stronger for PN functionalized with RNA sequences such as m*-RNA^Cy5^ (3), U_20_-RNA^Cy5^ (5) compared to PN simply functionalized with m*^Atto647N^ (2). However, other EMSA results are in stark contrast to the partitioning data. Pristine PN (1) and 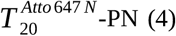 do not carry the strongly binding RNA sequence and yet show the highest partitioning values. Most interestingly, m/m*^Atto647N^ (2) shows high binding affinity in EMSA, but the lowest partitioning on the PN level. The highest partitioning can be found for 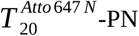 with pristine m barcode, but with hybridized A_20_/T_20_ sequences. Taken together, these experiments highlight for the first time the impact of environmental parameters on nucleic acid/FUS interactions. Most critically, standard in-solution EMSA affinity measurements deliver in parts completely different interaction maps compared to the crowded and DNA-rich PN environment. The most significant disparity can be identified for the comparison of 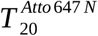 (4, P = 743) and m*^Atto647N^ (2, P = 36) with a ca. 20-fold difference in partitioning (Figure 4d), although both duplexes display similar EMSA affinities (Figure 4e, columns A_20_/T_20_ and m/m*).

### PN core modifications alter dynamics and condensate morphology

Next, we turn to kinetic details of FUS-GFP PS in these differently functionalized PN, after initiating PS by cleavage of the MBP-tag with TEV protease. Note that TEV cleavage is a rapid process, completed within less than 2 min in buffered solution (Supplementary Fig. 17). In pristine PN (1), PS starts also quickly, with FUS-GFP droplet formation occurring within five minutes after MBP cleavage and subsequent wetting of the inner surface of the PN shell after a few hours (Figure 4c, left). Additionally, the remaining droplets increase in size due to coalescence. Over the course of 1-2 days the wetting region gains in thickness due to coalescence of FUS-GFP droplets with the FUS-GFP shell. We further comment on the formation of the concentric shell below for dye-labeled PN.

Introduction of m*^Atto647N^ (2) leads to key differences in the PS behavior of FUS-GFP. Hardly any FUS-GFP droplets form in the initial 30 minutes after TEV cleavage within the PN. Afterwards droplets are formed inside and on the outside surface of the PN. Note, that m*^Atto647N^ (2) visualizes the PN core material (magenta channel) and shows a uniform distribution in the time series. Despite low partitioning and bad FUS-GFP retention (compare Supplementary Fig. 18 and 19), the final image shows colocalization of the FUS-GFP droplets and the core material (single channel images: Supplementary Fig. 9 and 10), indicating that p(A_20_-m/m*^Atto647N^) chains co-separate with the FUS-GFP condensates.

For the other samples, FUS-GFP PS progresses more slowly. This is not only due to chemical interactions, but also due to decreased core dynamics as identified by FRAP of all modified PN before MBP-FUS-GFP addition (Figure 4f). m*^Atto647N^-PN (2) show a FRAP recovery with *t*_1/2_ = 1.4 s (FRAP details in section 9 of the SI, Supplementary Fig. 13, 14), whereas, in comparison, all other modified PN do not show any appreciable recovery in the same time frame. The reason for the low dynamics in the RNA-containing samples is the fact that the dangling RNA strand is in fact a hairpin that can also dimerize at high concentrations inside the PN instead of only folding intramolecularly in dilute solution (Supplementary Table 1, also containing NUPACK secondary structure simulations of dimers). In terms of structure formation, in m*-RNA ^Cy5^-PN (3), the FUS-GFP condensates form in the center of the PN and do not show wetting of the inner surface of the shell as in pristine PN (1). In direct comparison to the m*^Atto647N^-PN (2), the addition of dangling RNA prevents leakage of the FUS-GFP condensates or formation of droplets at the outside of the shell. Overall, the retention of FUS-GFP after 5 h is much higher for the RNA-containing m*-RNA^Cy5^-PN (3) than for m*^Atto647N^-PN (2). The FUS-GFP condensates still behave dynamically as seen from the droplets merging from 5 h to 2 days, even though FUS-GFP and PN core material clearly form a combined phase. This observation is supported by FRAP measurements of the phase-separated droplets (Figure 4g), showing a partial recovery of FUS-GFP after 1 day, while the core channel remains non-dynamic. This means that FUS-GFP condensate droplets can exchange RNA partners in the core, allowing for diffusion inside the PN core.

The remaining samples 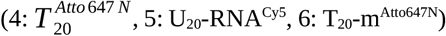 are dominated by slow PN core dynamics. In all cases the phase separated FUS-GFP forms interconnected networks within the PN, which is an unusual behavior for the minority phase. It is a prime example of viscoelastic phase separation (VPS) known from polymer physics, which is caused by a dynamic asymmetry of two components in a mixture, here the long ssDNA in the PN core and FUS-GFP.^57^ This dynamic asymmetry leads to the slower phase sustaining mechanical stress, which results in the formation of network-like morphologies instead of droplets. For 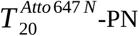 (4), the phase separated structure remains arrested within two days, whereas for U_20_-RNA^Cy5^ (5) and T_20_-m^Atto647N^ (6) PN, this VPS-network relaxes slowly over the span of two days. Here, the affinity of FUS-GFP to the PN material is strong enough to induce co-PS. This leads to vacuole formation, with the condensate being segregated to the PN shell. The formation of the concentric shell is due to affinity of the pA core material with the pT shell (A_20_/T_20_-duplexes stabilize the PN core shell interface; see Scheme 1a). The similarity in the final morphology compared to pristine PN (1) raises the question whether the pA core material of the pristine PN is segregated as well. Indeed, a covalently labelled p(A_20_-m)^Atto425^ shows a similar co-separation (Supplementary Fig. 20), which is also why we selected parameters in the numerical simulations (Figure 3) to reflect this behavior. At first glance, this A_20_/T_20_-duplex driven morphology seems unique to PN, yet it bears some resemblance to nuclear organization in cells. Cell nuclei are organized by spatially segregated euchromatin and heterochromatin domains, with the latter located at the nuclear periphery. This structure results from tethering of heterochromatin to nuclear lamina, which are rooted in the inner nuclear membrane.^58,59^ PN abstract this concentric organization in a minimalist model system.

In summary, FUS-GFP partitioning, PS timescale and condensate morphology are drastically influenced by the nucleic acid sequences present in the PN core. Simple sequence variations lead to the extremes of slowed down PS kinetics 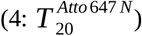, partial expulsion of FUS-GFP from the core (2: m*^Atto647N^) and full co-PS of FUS-GFP with the PN core material and segregation to the interface with vacuole formation (1: pristine, 5: U_20_-RNA^Cy5^, 6: T_20_-m^Atto647N^). Critically, classical EMSA affinity measurements largely fail to predict the partitioning behavior observed in our complex system. This disparity underscores a central insight of our study: Protein/nucleic acid affinity alone cannot predict polymer behavior in confined, crowded, and multivalently interacting PN environments. Instead, the dynamic and structural context in which these interactions occur plays a decisive role. The origin relates to the fact that the PN permits multimodal binding in a compact, electrostatically diverse space, instead of an isolated binary interaction. For instance, high affinity binders like m*-RNA^Cy5^ (3, Figure 4e) cause strong multimodal binding which restricts the degrees of freedom of the protein by locking its orientation. In contrast to this restricted conformation, a weak binder will offer conformational freedom to the protein, allowing it to more dynamically interact with an environment phase and adapt to its constraints. We hypothesize that these weak, transient interactions provide a better environment for partitioning in crowded systems. While our results indicate intricate and complex interplays between many parameters, they clearly show that test-tube affinity measurements have limitations in replicating the processes in crowded DNA-rich environments, highlighting the importance of developing PN to better mimic intra-nuclear complexity.

### FUS PS is controlled by PN core crosslinking

Next to tuning the composition, another key advantage of our PN platform is the ability to tune their dynamic and viscoelastic properties. This can be achieved by using a modified p(A_20_-m-XL) sequence, which contains a self-complementary region XL that forms crosslinks (Figure 5 inset). Such linkers are known to profoundly impact the viscoelastic properties, as well-established for macroscopic DNA hydrogels.^60,61^ Mixing p(A_20_-m-XL) with p(A_20_-m) prior to the formation of the PN leads to co-assembly within the PN, allowing to fine-tune their composition. FRAP measurements after MBP-FUS-GFP partitioning reveal a great impact of the XL domain on PN dynamics (Supplementary Fig. 21). The characteristic *t*_1/2_ increases for higher p(A_20_-m-XL) fractions and the endpoint recovery (ER) decreases, confirming the intended decrease in PN core dynamics for higher XL amounts. Complementary to this quantitative mobility analysis, Figure 5 shows a morphology progression over time. Together, these datasets offer a comparative analysis of the FUS PS dynamics upon PN core crosslinking.

**Figure 5:**
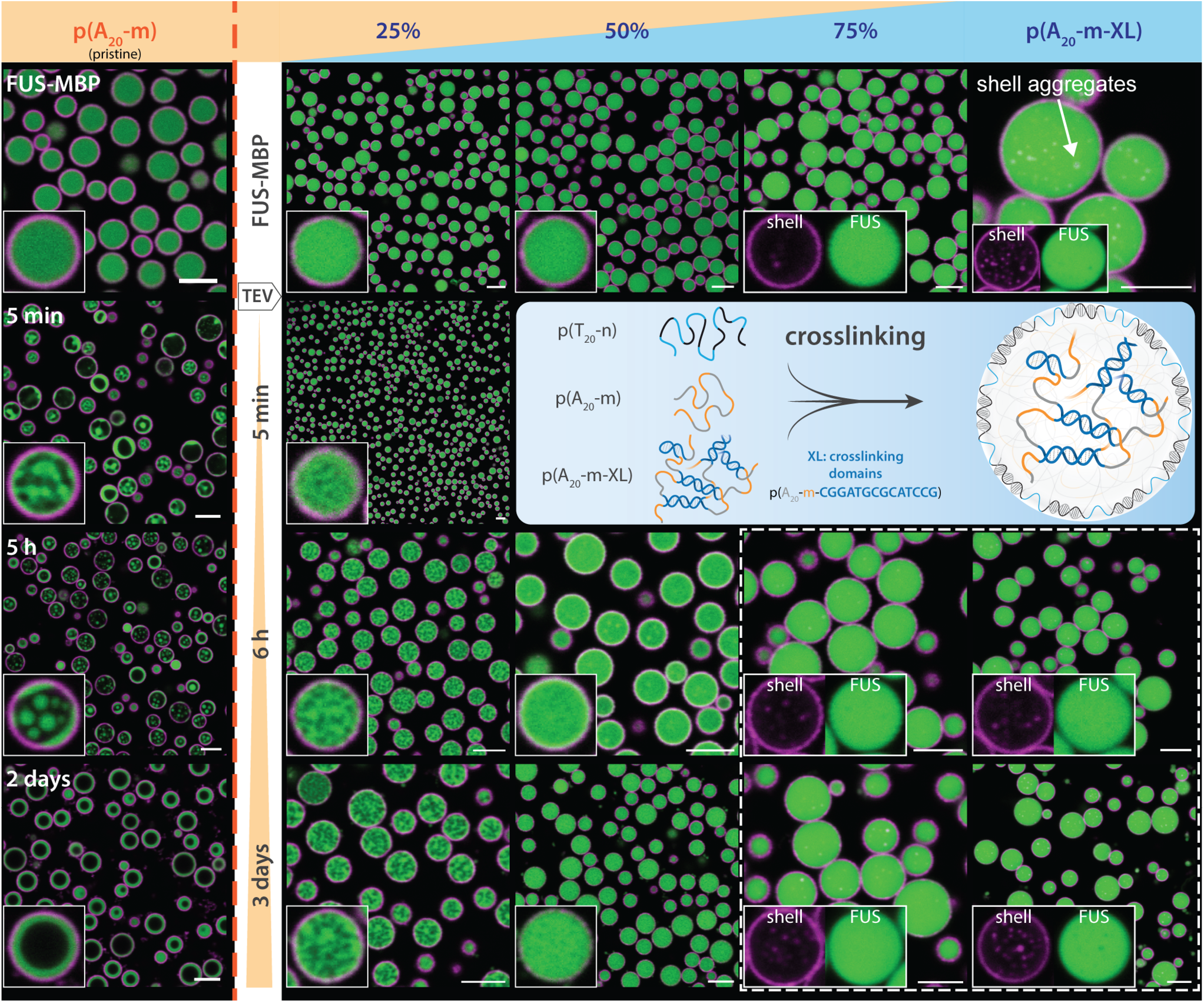
Crosslinking of PN influences FUS PS. Various degrees of crosslinking can be achieved by adding different fractions of p(A_20_-m-XL) to the PN (LR = 0.30). XL forms self-complementary crosslinks. Both highly crosslinked samples (75% and 100%) show some magenta speckles in the PN core, which originate from some shell aggregates present before TEV addition. The dashed square in the bottom right indicates a region of no visible PS within the optical limits of the microscope. A corresponding FRAP experiment is shown in Supplementary Fig. 21. Scale bars: 5 µm.

A clear trend emerges how crosslinking affects FUS PS in PN after addition of TEV protease. Non-crosslinked, pristine PN with pure p(A_20_-m) display quick FUS PS and droplet formation that reorganizes over the course of 2 days to a concentric shell (same data as shown in Figure 4c). These dynamics are greatly reduced when adding 25% p(A_20_-m-XL), which changes both the PS morphology and time progression. Grainy FUS PS is visible after 5 min, which only slightly coarsens during the next 3 days. Instead of localizing at the inner shell, a spongy and arrested phase-separated network occurs that originates from VPS.^57^ The basis for this behavior lies in the dynamic asymmetry created by the slower dynamics of the crosslinked core, as already discussed above (Figure 4). The XL motifs impose additional connectivity within the PN core, thereby increasing the energetic and kinetic cost of redistributing the DNA core during FUS condensation. This restricts domain growth and relaxation, so that the 25% crosslinked sample remains trapped in a finely structured, spongy morphology rather than progressing toward larger, coarsened FUS-rich phases. A further increase of crosslinker leads to even slower dynamics. For 50% p(A_20_-m-XL), FUS PS is barely visible after 6 h and does not progress much over the next 3 days. Finally, at 75% crosslinking and above, no microscopically visible FUS PS occurs anymore over the span of 3 days.

### Viscoelastic environment suppresses FUS liquid-to-solid transitions

As commonly reported for FUS, condensates can age and undergo a liquid-to-solid transition over time.^23,24,62–64^ Our PN with tunable dynamics can help to understand how confinement can influence this behavior. FUS condensate liquid-to-solid transition is measurable by FRAP of the protein in buffer (droplet assay).^65,66^ Since FUS only shows PS after MBP cleavage, there is no direct reference sample for FRAP of the fully dissolved MBP-FUS-GFP state. As a substitution for this data point, we prepared a sample with 10% PEG as a molecular crowder which forces MBP-FUS-GFP PS even without MBP cleavage (Figure 6a). The remaining data points after TEV addition are taken from a droplet assay without PEG and in presence of the low-molecular PN core sequence (A_20_-m)_5_ with m*-RNA^Cy5^. DNA and RNA have been added in the same amounts as for PN samples, with the difference of not being assembled into PN (see SI section 16; Supplementary Fig. 22). As expected, these samples show a strong decrease in FUS mobility already after 6 h in buffer on account of ongoing aging (Figure 6a).

**Figure 6:**
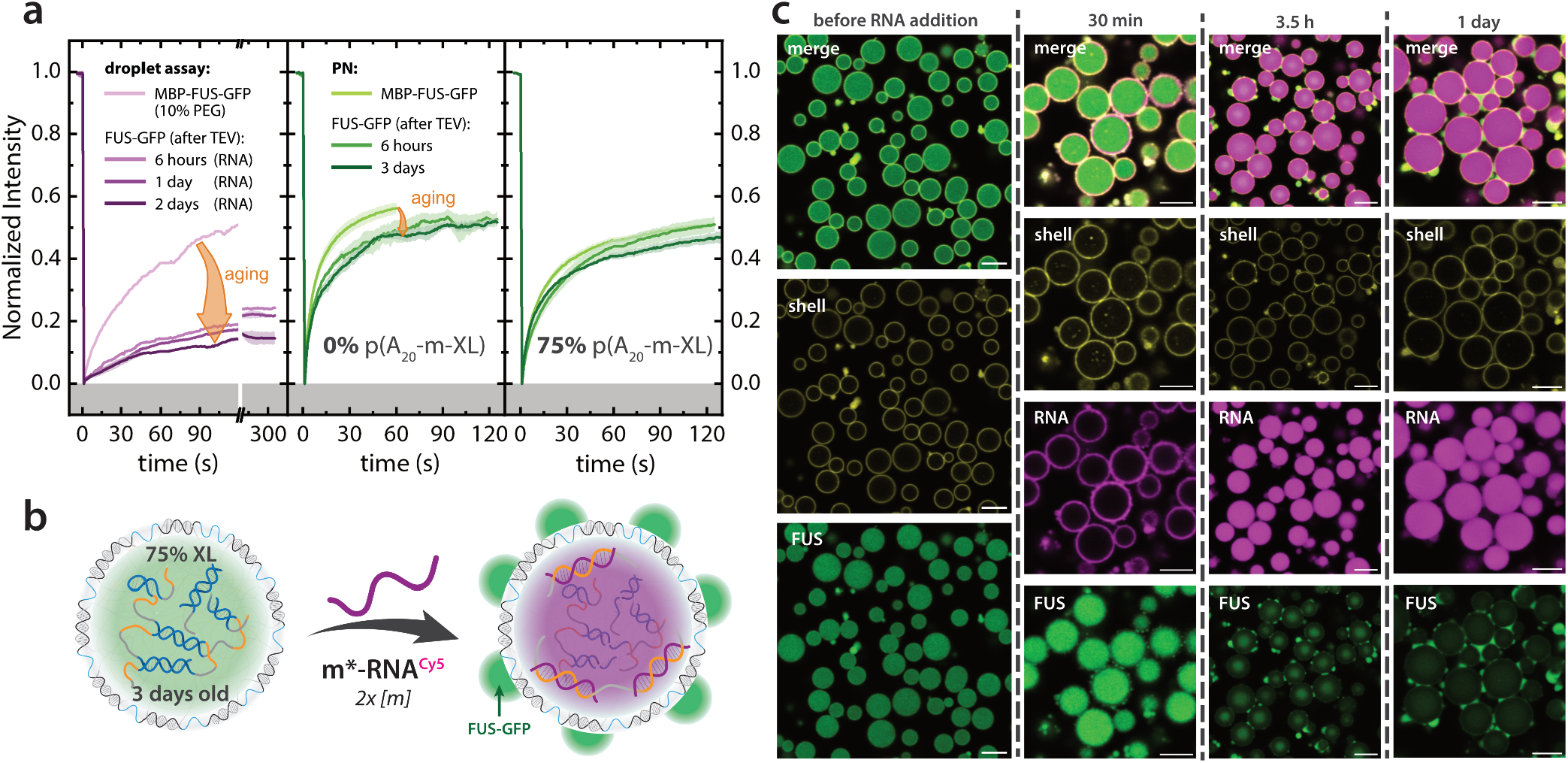
FUS-GFP in PN remains dynamic after 3 days. **a** FRAP measurements in the FUS-GFP channel in a droplet assay (left) versus 0% and 75% crosslinked PN samples (right) at different time points: After MBP-FUS-GFP addition (light purple and green line), and at different time points after TEV protease induced PS (darker purple and green lines). Droplet assay: N = 1-3. PN 0% p(A _20_-m-XL): N = 3-5. PN 75% p(A_20_-m-XL): N = 3-5. N denotes the number of individual FRAP measurements. **b** scheme and **c** CLSM images showing FUS-GFP is drawn out of 75% XL-PN after addition of excess m*-RNA^Cy5^ to a 3 days old PN sample. Scale bars = 5 µm.

Having established these reference systems, we turn to the PN systems with FUS-GFP. The pristine p(A_20_-m)-PN without crosslinking show only a small consecutive decrease in the FRAP recovery profiles for FUS-GFP over time, whereas a clear decrease to before PS (MBP-FUS-GFP) is discernible (Figure 6a). For PN with p(A_20_-m-XL)/p(A_20_-m) = 75/25, hence at highly crosslinked state, the initial FRAP of FUS-GFP shows lower recovery dynamics due to the highly crosslinked nature of the PN. However, as time progresses, a further liquid-to-solid transition of FUS condensates is suppressed, as the decrease of the FRAP profiles is comparably little in highly cross-linked PN, as well as on the same level as to before PS (MBP-FUS-GFP) (Figure 6a). This led us to conclude that both PS and aging of phase-separated FUS are suppressed by crosslinking.

To further underscore the dynamic properties of FUS-GFP in the 75% crosslinked PN, we added a two-fold excess of m*-RNA^Cy5^ to offer FUS-GFP a non-polymeric binder after 3 days of aging inside the PN (Figure 6b). This control experiment is designed to discern liquid FUS from solid fractions, as we hypothesized that non-solidified FUS would interact and phase-separate with the dynamic nucleic acid binder. Over the course of a few hours, the m*-RNA^Cy5^ infiltrates the core of the aged, 75% crosslinked PN (Figure 6c). However, because of the two-fold excess there is a large unbound fraction of m*-RNA^Cy5^ in solution, so that FUS-GFP condensates form at the PN–solution interface with only a low intensity in the green FUS channel remaining inside the PN. We suggest that the residual FUS-GFP in the PN core largely originates from the solid fraction, whereas the dynamic, RNA-bound fraction is leached out and forms condensates on the outer surface of the PN shell.

Taken together, the viscoelastic suppression of FUS-GFP PS also slows down the aberrant liquid-to-solid transition of the protein – a hallmark of disease. Hence, the viscoelastic properties are not only relevant to enter the domain of VPS (FUS network formation, although at binodal concentrations), but also to regulate the extent of FUS PS and its liquid-to-solid transition during aging. These results are instructive for possible therapeutic strategies, as they highlight the importance of the physical environment of membraneless organelles for aberrant phase transition and protein aggregation processes.

## Conclusions

This work has introduced protonuclei (PN) – programmable DNA-based nucleus-inspired compartments – as a versatile platform to study the impact of environmental constraints on the affinity, phase separation (PS), and liquid-to-solid phase transitions of nucleic acid-binding proteins. Our findings give a new perspective on how to interpret protein-nucleic acid interactions and PS in the context of more complex cell-like environments, here in particular the cell nucleus. While affinity-based assays remain useful, they fail to reflect the complex interplay of multivalency, confinement, and physical state that governs condensation in the nucleus. The PN system and future derivates thereof have the potential to fill this gap, offering a controllable testbed that reveals how local context shapes condensate formation, retention, and aging. By mimicking essential features of the cell nucleus, including nucleic acid crowding, compositional heterogeneity, and tunable viscoelasticity, PN provide a new intermediate between simplistic *ex vivo* test tube assays and complex cellular systems. Using FUS, a protein linked to neurodegenerative diseases, we demonstrate that its phase behavior inside PN is strongly influenced by the sequence and structure of nucleic acids, as well as by the dynamic mechanical properties of the DNA-rich core.

A central finding is the limited predictive power of solution-based protein-nucleic acid interaction assays, such as EMSA, which fail to capture the complex interactions arising in the multivalent and crowded PN environment. We show here that seemingly minor changes in nucleic acid sequence or mechanical constraints can have drastic effects on protein partitioning, phase morphology, and condensate dynamics. Moreover, by incorporating crosslinks into the PN core, we demonstrate how viscoelasticity can suppress PS and delay the liquid-to-solid transition during condensate aging – pointing to a critical role of the physical environment in regulating phase behavior. The ability of PN to suppress PS or alter condensate morphology via crosslinking parallels how nuclear viscoelasticity regulates biological function *in vivo*. For instance, mechanical deformation during confined cell migration induces chromatin reorganization that can trigger condensate coalescence and de novo assembly.^67^ Furthermore, nuclear stiffness is dynamically tuned by chromatin compaction, which suppresses PS in stiff heterochromatin regions and allows PS in softer euchromatin regions. By using targeted condensation and stiffness gradients, it is assumed that cell nuclei can dynamically restructure their genomic loci. ^68^ This modulation of nuclear PS in cells points to potential physical regulators of phase transitions beyond biochemical interactions. These insights support emerging views that pathological phase transitions may be mitigated not only by molecular inhibitors but also by targeting the mechanical properties of the nuclear environment.^69^

Looking ahead, the PN system in combination with numerical simulations, offers a powerful framework to probe more complex nuclear PS phenomena. Its modularity allows incorporation of multiple proteins and nucleic acids, enabling studies of competition, cooperativity, and emergent behavior. The ability to systematically vary protein variants and nucleic acid sequences also opens up new routes to explore disease-associated mutations and their impact on phase transitions in a nucleus-like environment. Additionally, future development of a theoretical model to predict protein PS in PN can be an insightful tool to identify the underlying molecular interactions governing partitioning, dynamics, morphology and PS suppression. Beyond structural observations, PN can be adapted to investigate the functional consequences of PS, for example on transcription or RNA processing. Given their synthetic nature and programmability, PN may also serve as bottom-up tools for screening condensate-modulating compounds, or as functional components in synthetic biology systems.

In the long term, the PN platform could aid in reconstituting complex, disease-relevant membraneless organelles under tunable conditions. This might allow systematic dissection of how disease-linked mutations, co-factors, or mechanical perturbations influence phase behavior – a task that remains challenging in live cells. As synthetic biology and condensate pharmacology advance, synthetic systems like PN may also help screen for compounds that modulate phase behavior not just by disrupting binding, but by altering the physical microenvironment. Altogether, our findings not only highlight the limitations of classical assays but also provide a blueprint for powerful *ex vivo* models. PN offer a rich design space to dissect the rules of intracellular PS and hold potential for guiding future therapeutic strategies targeting condensate dynamics in disease.^70,71^

## Methods

### Materials

Oligonucleotide strands were ordered as shown in Supplementary Table 1 from Integrated DNA Technologies (IDT) and Biomers. Enzymes phi29 DNA Polymerase (10 U μL^−1^), T4 DNA Ligase (2 U μL^−1^), Exonuclease I (20 U μL^−1^), Exonuclease III (200 U μL^−1^) were purchased from LGC Genomics. Thermostable inorganic pyrophosphatase (TIPP, 2000 U μL^−1^) and nuclease-free water were purchased from New England Biolabs (NEB), dNTPs (100 mM - 110 mM) were purchased from Jena Bioscience. Sodium chloride (Gen-Apex) was purchased from VWR, magnesium chloride solution (2M, BioUltra) was purchased from Sigma Aldrich, Invitrogen TE buffer (10 mM Tris, 1 mM EDTA, pH 8.0) was purchased from Thermo Fisher Scientific.

### PN core and shell ssDNA synthesis

The procedure for rolling circle amplification of ssDNA polymers was adapted from our previous report.^28^ The stock solutions of 5′-phosphorylated templates and their corresponding ligation strand (Supplementary Table 1) were mixed in equimolar stoichiometry to attain a final concentration of 1 μM in TE buffer (Invitrogen; 10 mM Tris(hydroxymethyl)aminomethane pH = 8 and 1 mM EDTA) containing additionally 100 mM NaCl (total volume 100 μL). The concentrations of the strands are kept low to prevent inter-strand hybridizations. The buffered mixture was heated up to 85 °C (for 5 min) at a rate of 3 °C s^−1^ and slowly cooled down to 25 °C at 0.01 °C s^−1^. After annealing the strands, 20 μL of 10x commercial ligase buffer (LGC; 500 mM Tris-HCI, 100 mM MgCl_2_, 50 mM dithiothreitol, and 10 mM ATP), 70 μL of water, and 10 μL of T4 ligase (2 U μL^−1^) were added in the reaction tube containing 100 μL of the template strand, stirred (10 min, 400 rpm) and left to react for 3 h at 25 °C. The T4 ligase was then denatured by heating the reaction mixture for 20 min at 70 °C. Then 10 μL of exonuclease I (LGC; 40 U μL^−1^) and 10 μL of Exonuclease III (LGC; 200 U μL^−1^) were added, and the mixture was kept overnight at 37 °C on a thermo-shaker (Eppendorf) with gentle shaking (300 rpm) to digest unreacted ligation strands and non-circularized templates in solution. Then the exonucleases were subsequently deactivated by heating the reaction mixture at 80 °C for 40 min. The circular templates were purified using the Amicon Ultra-centrifugal filters with a 10 kDa cut-off (Merck Millipore) and washed three times using TE buffer over the same filter. The concentration of circular templates was measured by the use of a ScanDrop (Jena Analytic) spectrophotometer, and the solutions were diluted to 10 μM using TE buffer. To synthesize multiblock ssDNA polymers, 0.5 μL circular template (10 μM) were mixed with 10 μL of commercial 10x polymerase buffer (LGC; 500 mM Tris-HCl, 100 mM (NH_4_)_2_SO_4_, 40 mM Dithiothreitol, 100 mM MgCl_2_), 1 μL of exonuclease resistant primer (10 μM in TE buffer), 2 μL of Φ29 Polymerase (LGC; 10 U μL^−1^), 5 μL of pyrophosphatase (New England Biolabs; 2000 U μL^−1^), and 5 μL of an adjusted dNTP mixture (total dNTP concentration of 5 mM, the percentage of each base correspond to the expected sequence composition in the ssDNA polymer), and 76.5 μL of ultrapure nuclease-free water. The reaction mixtures were kept at 30 °C for 60 h on a thermo-shaker with a gentle shaking (300 rpm). The solution containing ultralong ssDNA polymer was subjected to temperature-induced cleavage for 15 min at 95 °C, and the resulting solution was then washed through Amicon Ultra-centrifugal filters with a 30 kDa cut-off (Merck Millipore, three to four times) using 350 μL of TE buffer. The amount and purity of the ssDNA polymers were determined using a ScanDrop (Jena Analytic) spectrophotometer. The pyrophosphatase enzyme was added to prevent the formation of insoluble magnesium pyrophosphate as a side product that hinders the polymerization process and is also accountable for resulting in so-called nanoflowers and might cause misperception during the microscopic investigations.

### Purification of MBP-FUS-EGFP-His_6_ and MBP-FUS-His_6_

Expression and purification of recombinant MBP-FUS-EGFP-His_6_ and MBP-FUS-His_6_ was performed as described in (Hofweber, Cell 2018)^64^. Briefly, BL21-DE3-Rosetta-LysS cells were transformed with a pMal-MBP-Flag-FUS-GFP-His_6_ or MBP-FUS-His_6_ expression plasmid and grown in standard lysogeny broth (LB) medium until OD (600 nm) value reached 0.8. Expression was induced with 0.1 mM IPTG for 22 h at 12 °C. Cells were lysed in lysis buffer (50 mM Na_2_HPO_4_/NaH_2_PO_4_ pH 8.0, 300 mM NaCl, 10 μM ZnSO_4_, 40 mM imidazole, 4 mM β-mercaptoethanol, 10 % (v/v) glycerol and 1 μg/ml each aprotinin, leupeptin and pepstatin) using sonication. MBP-FUS-GFP-His_6_ or MBP-FUS-His_6_ were purified by tandem-affinity purification using Ni-NTA agarose (Qiagen) and amylose resin (NEB) and eluted in lysis buffer supplemented with 250 mM imidazole and 20 mM maltose for MBP-FUS-EGFP-His_6_, and 50 mM imidazole and 10 mM maltose for MBP-FUS-His_6_, respectively. Proteins were dialyzed against storage buffer (20 mM Na_2_HPO_4_/NaH_2_PO_4_, pH 8.0, 150 mM NaCl, 5% (v/v) glycerol, 1 mM EDTA, 1 mM DTT), snap-frozen in liquid N_2_ and stored at -80°C. The OD260/280 ratio for MBP-FUS-EGFP-His_6_ was 0.74, and for MBP-FUS-His_6_ was 0.65.

### His_6_-TEV protease purification

Expression and purification of His_6_-Tev was performed as described in (Hofweber, Cell 2018)^64^. Briefly, BL21-DE3 Rosetta-LysS cells were transformed with a pET24(+)-His-Tev plasmid and grown in standard lysogeny broth (LB) medium until OD (600 nm) value reached 0.7. Expression was induced with 0.5 mM IPTG for 20 h at 18 °C. Cells were lysed in lysis buffer (50 mM Tris pH8, 200 mM NaCl, 20 mM imidazole, 10% (v/v) glycerol, 4 mM β-mercaptoethanol) in presence of 0.1 mg/ml RNase A using lysozyme and sonication. His_6_-Tev was purified using Ni-NTA agarose (QIAGEN) and eluted in lysis buffer containing 600 mM imidazole. The protein was dialyzed against storage buffer (50 mM Tris pH8, 150 mM NaCl, 20% (v/v) glycerol, 2 mM DTT), snap-frozen in liquid N_2_ and stored at -80°C.

### PN formation

PN are formed as shown in Scheme 1a. The formation of PN is initiated by combining pA at 0.18 g/L and pT at 0.03 g/L in TE buffer, together with MgCl_2_ at a final concentration of 50 mM. A typical preparation would target a final volume of 65 μL in a PCR tube, which is enough for 6 experimental batches. The mixture is then subjected to a heating ramp to 95°C for 20 min with a heating and cooling rate of 3°Cmin^−1^. After cooling the sample is split up and 10 μL of the PN stock are added to a fresh PCR tube containing the respective dye-labelled DNA or RNA strands. Here, the DNA complementary to the shell (n*) is added in exact stoichiometry to the barcode concentration, whereas the core DNA or RNA (m*) is added with a dilution factor of 0.8 to ensure there is none left in solution. After mixing with a pipette, the samples are allowed to equilibrate for 60 minutes while shaking at 400 rpm. In a final step the buffer is adjusted to FUS-nucleic acid binding buffer by 10x dilution with HEPES buffer (10 mM HEPES, 200 mM KCl, 1 mM DTT, 5 mM MgCl_2_, Glycerol 3%, pH 7.5) in a well plate, yielding a final Mg^2+^ concentration of 10 mM. The PN are then allowed to swell and settle to the bottom of the plate, so that the volume can be carefully reduced with a pipette from the top to a working volume of 25 μL. During this process the plate is sealed to prevent evaporation, the seal is only lifted for further pipetting steps, e.g. TEV addition. Plates are generally stored at room temperature (25 °C) during all steps.

For crosslinked PN the specified fraction of pA was replaced by pA-XL with a self-complementary domain, but all other steps of the preparation remain the same.

### Concentration-dependent PS experiments

PN were prepared as mentioned above, but without the addition of core-modifying DNA/RNA strands. In order to stay within the same PN batch for all experiments, half of the PN stock was stored at 4 °C for three days prior to use. We routinely observe PN stocks in high-salt buffer to be stable for weeks to months, highlighting their exceptional robustness. Imaging of the samples was done continuously for the first 30 min after TEV addition and again after 5 h. All images have been adjusted in contrast and brightness.

### PN core modification experiments and crosslinking

A fresh aliquot of the MBP-FUS-GFP protein (45 μM) was thawed slowly on ice and mixed gently with a pipette. It was then added at a final concentration of 1.5 μM to the respective PN microplate well with a pipette and immediately mixed with a bigger volume pipette to ensure an even distribution. The protein was allowed to partition into the PN for 30-60 min at room temperature until completion of partitioning was confirmed via CLSM. Then a freshly thawed aliquot of TEV protease (80 μM) was mixed and added to the wells at a final concentration of 5 μM, marking the start of the PS process. After all additions were made, the plate was sealed to avoid evaporation within the timespan of the experiment.

### Electrophoretic Mobility Shift Assay (EMSA)

The nucleic acid sequences, modifications and suppliers can be found in Supplementary Table 1. Duplexes (m/m*^Atto647N^ and 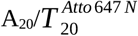) were annealed using one labelled strand and one unlabelled strand. 500 nM of labelled nucleotide probe were diluted in binding buffer (20 mM Na-Phosphate pH 7.5, 75 mM NaCl, 5 mM MgCl _2_, 2.5 % Glycerol,1 mM DTT, 1 U/µl RNase inhibitor (Thermo Fisher)) and mixed with varying amounts of MBP-FUS-His_6_. Binding reactions (10 µl) were incubated for 1 h at RT and after addition of 2 µl 30% (v/v) glycerol samples were loaded onto a 1 mm thick non-denaturing polyacrylamide gel (6%) in 0.5 x TBE. Gels were run at 100 V for 40 min at RT before imaging with a Bio-Rad ChemiDoc MP Imaging system.

## Supporting information

complete SI

## Acknowledgements

We acknowledge funding from CRC1551 ‘Polymer Concepts in Cellular Function’ of the Deutsche Forschungsgemeinschaft (DFG project number 464588647). A.S. acknowledges support through a postdoctoral grant of the Alexander von Humboldt Foundation. D.D. acknowledges funding by the DFG Heisenberg programme (DFG project number 442698351). E.S. was supported by a Peter and Traudl Engelhorn Postdoctoral Fellowship. Parts of this research were conducted using the supercomputer MOGON 2 and/or advisory services offered by Johannes Gutenberg University Mainz (hpc.uni-mainz.de), which is a member of the AHRP (Alliance for High Performance Computing in Rhineland Palatinate, www.ahrp.info) and the Gauss Alliance e.V. The authors gratefully acknowledge the computing time granted on the supercomputer MOGON 2 at Johannes Gutenberg University Mainz (hpc.uni-mainz.de). We would like to thank Prof. Dr. Miguel Andrade for his helpful discussions and Sára Varga for the provision of additional purified TEV and MBP.

## Author Contributions

D.D. and A.W. conceived and supervised the project. E.S. expressed and purified the protein. N.K. performed the EMSA experiments. J.F. planned and carried out all other experiments, performed data analysis, and was responsible for the majority of the experimental work. Numerical simulations were conducted by J.J.M., and all-atom MD simulations were performed by A.P. under the supervision of L.S. Initial experiments and early polymer synthesis were performed by A.S.. J.F. and A.W. wrote the manuscript, with input from all authors.

## Abbreviations

ALS: amyotrophic lateral sclerosis
CLSM: confocal laser scanning microscopy
DNA: deoxyribonucleic acid
eGFP: enhanced green fluorescent protein
EMSA: electrophoretic mobility shift assay
ER: endpoint recovery
FRAP: fluorescence recovery after photobleaching FTD frontotemporal dementia
FUS: fused in sarcoma protein
IDR: intrinsically disordered region
MBP: maltose binding protein
NLS: nuclear localization signal
PN: protonucleus / protonuclei
PS: phase separation
RNA: ribonucleic acid
TEV: Tobacco Etch Virus protease
TNPO1: Transportin-1
VPS: viscoelastic phase separation

