## Supplementary material for "Programmable DNA Protonuclei Reveal Environmental Context on Protein Phase Separation": complete SI

#### Supplementary Information

Johann Fritzen[1], Avik Samanta[2], Nele S. Kuhr[3], Antonia Preuß[3,4], Erin L. Sternburg[3], Lukas S. Stelzl[3,4], Jasper J. Michels[5*], Dorothee Dormann[3,4]*, Andreas Walther[1,5]*

[1] Life-Like Materials and Systems, Johannes Gutenberg University, 55128 Mainz, Germany
[2] Indian Institute of Technology Kharagpur, West Bengal, India
[3] Institute of Molecular Physiology, Johannes Gutenberg University, 55128 Mainz, Germany
[4] Institute of Molecular Biology (IMB), 55128 Mainz, Germany.
[5] Max Planck Institute for Polymer Research, 55128 Mainz, Germany

### 1. DNA & RNA sequences

**Supplementary Table 1: Nucleic acid sequences and modifications.** DNA sequences are given as capital letters, whereas RNA sequences use lowercase letters. An asterix (*) between two bases indicates a bridging phosphorothioate modification (PTO) at the designated position. An asterix in the sequence's name indicates it is complimentary to the specified domain, e.g. m* being complimentary to m.

| | Name | Sequence (5'→3') | Modification | Supplier |
|---|---|---|---|---|
| *core DNA p(A$_{20}$-m)* | Template | ATC TAT CCT AAT TTT TTT TTT TTT TTT TTT TGA ACC CGT AT | 5'-Phosphorylation | IDT |
| | Ligation | TTA GGA TAG ATA TAC GGG TTC | - | IDT |
| | Primer | TTA GGA TAG ATA TAC GGG T*T*C | PTO | IDT |
| | Polymer | [AAA AAA AAA AAA AAA AAA AAT TAG GAT AGA TAT ACG GGT TC]$_x$ | - | RCA |
| *shell DNA p(T$_{20}$-n)* | Template | ATC CTC TAA AAT CAA AAA AAA AAA AAA AAA AAA GTA AAA CCA CAC G | 5'-Phosphorylation | IDT |
| | Ligation | TTT TAG AGG ATC GTG TGG TTT T | - | IDT |
| | Primer | TTT TAG AGG ATC GTG TGG TT*T* T | PTO | IDT |
| | Polymer | [TTT TTT TTT TTT TTT TTT TTG ATT TTA GAG GAT CGT GTG GTT TTA C]$_y$ | - | RCA |
| *core DNA p(A$_{20}$-m-XL)* | Template | ATC TAT CCT AAT TTT TTT TTT TTT TTT TTT TCG GAT GCG CAT CCG GAA CCC GTA T | 5'-Phosphorylation | IDT |
| | Ligation | TTA GGA TAG ATA TAC GGG TTC | - | IDT |
| | Primer | TTA GGA TAG ATA TAC GGG T*T*C | PTO | IDT |
| | Polymer | [AAA AAA AAA AAA AAA AAA AAT TAG GAT AGA TAT ACG GGT TCC GGA TGC GCA TCC G]$_x$ | - | RCA |
| *core modifications* | m*$^{Atto647N}$ | TGA ACC CGT ATA TCT ATC CTA A | 5'-Atto 647N | Biomers |
| | m*-RNA$^{Cy5}$ | ga acc cgu aua ucu auc cua agg auc uuu aac uac uca aga uac uga aca uga cau ggu a<br><br>*this RNA forms hairpin structures:*<br>Nupack MFE proxy structure at 25°C: [a)]<br>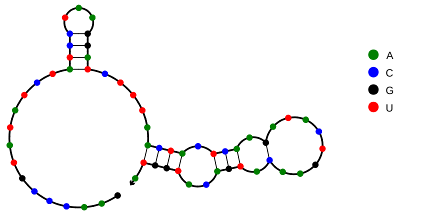<br><br>Free energy of the secondary structure: -6.65 kcal/mol<br><br>*dimerization is possible:*<br>Nupack MFE proxy structure at 25°C: [b)]<br>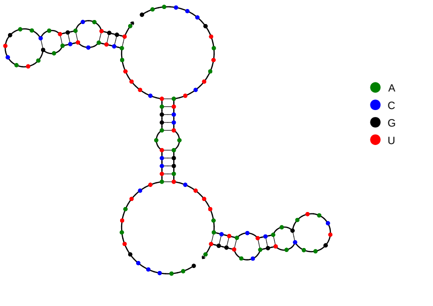<br><br>Free energy of the secondary structure: -25.13 kcal/mol | 5'-Cy5 | Biomers |

| | | | | |
|---|---|---|---|---|
| | $T_{20}^{Atto\,647\,N}$ | TTT TTT TTT TTT TTT TTT TT | 5’-Atto 647N | Biomers |
| | $U_{20}$-RNA$^{Cy5}$ | uuu uuu uuu uuu uuu uuu uug gau cuu uaa cua cuc aag aua cug aac aug aca ugg ua<br><br>*this RNA forms hairpin structures:*<br>Nupack MFE proxy structure at 25°C: [a)]<br>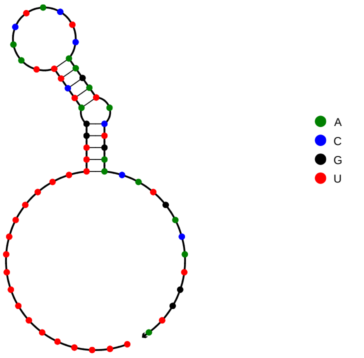<br><br>Free energy of the secondary structure: -6.27 kcal/mol<br><br>*dimerization is possible:*<br>Nupack MFE proxy structure at 25°C: [b)]<br>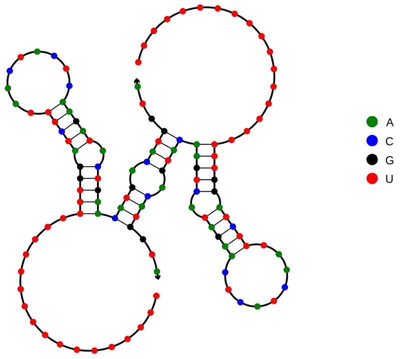<br><br>Free energy of the secondary structure: -26.10 kcal/mol | 5’-Cy5 | Biomers |
| | $T_{20}$-m$^{Atto647N}$ | TTT TTT TTT TTT TTT TTT TTT TAG GAT AGA TAT ACG GGT TC | 5’-Atto 647N | Biomers |
| *shell label* | n*$^{Atto565}$ | AAA ACC ACA CGA TCC TCT AAA A | 5’-Atto 565 | Biomers |
| *Strands for EMSA* | m | TTA GGA TAG ATA TAC GGG TTC | - | Biomers |
| | $A_{20}$ | AAA AAA AAA AAA AAA AAA AA | - | Biomers |

[a)] *Calculated with RNA model preset at 25°C, [RNA]=1 µM, max complex size 1.*
[b)] *Calculated with RNA model preset at 25°C, [RNA]=1 µM, max complex size 2.*

#### 2. Nucleic acid concentration in cell nucleus and in PN

The average diameter of a human cell nucleus is 6 µm[1]. Assuming spherical geometry, this amounts to a volume of 113.1 µm³. The size of the human genome is 3.2 billion base pairs, meaning 6.4 billion nucleotides. Consequently, human cell nuclei have a *c*(nt) of:

$$c(nt)_{upper}^{lit} = \frac{6.4 \times 10^{9} \ \ nt}{113.1 \ \ \mu m^{3}} \ = \ 56.59 \times 10^{6} \frac{nt}{\mu m^{3}} \tag{1}$$

Other reports mention up to 374 µm³ as a nuclear volume[2], which gives a lower limit of:

$$c(nt)_{lower}^{lit} = \frac{6.4 \times 10^{9} \ \ nt}{374 \ \ \mu m^{3}} \ = \ 17.11 \times 10^{6} \frac{nt}{\mu m^{3}} \tag{2}$$

The following formula is used to convert the values to g/L:

$$c(nt)^{lit}\left[\frac{g}{L}\right] = \frac{c(nt)^{lit}\left[\frac{nt}{\mu m^{3}}\right]}{6.022 * 10^{23} mol^{-1}} \times 330 \frac{g}{mol} \times 10^{15} \tag{3}$$

**Supplementary Table 2: Nucleotide concentrations in cells and protonuclei.**

| Specimen | Nucleotide concentration (nt/µm³) | Nucleotide concentration (g/L) |
|---|---|---|
| *Human cells, lower limit* | $17.11 \times 10^{6}$ | 9.4 |
| *Human cells, upper limit* | $56.59 \times 10^{6}$ | 31.0 |
| *PN, pristine* | - | 5.4 |
| *PN, 60 nt RNA* | - | 13.3 |

#### 3. Influence of TEV and MBP on PN morphology

The following control experiments show the influence of TEV, isolated MBP, and their combination on the morphology of covalently labelled PN core (magenta, dUTP-Atto425) and duplex-labelled shell (yellow, n*[Atto565]). We have followed the process at time points of 5 min, 30 min, 5 h, 1 day and 2 days for all three experiments. For all control experiments there are no changes to the PN morphology or PS of the core material, confirming that the auxiliary protein components (MBP solubility tag) and protease (TEV) remain innocent and FUS is the decisive protein to introduce PS.

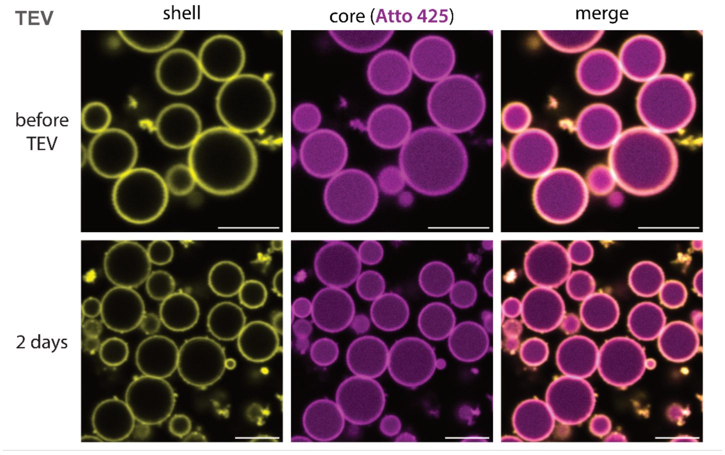

**Supplementary Fig. 1: Addition of TEV protease to pristine PN with a covalently labelled core does not induce PS.** Scale bars = 5 µm.

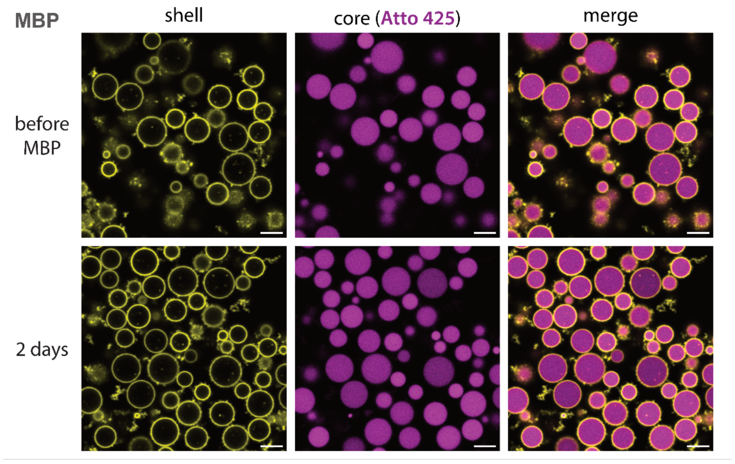

**Supplementary Fig. 2: Addition of isolated MBP to pristine PN with a covalently labelled core does not induce PS.** Scale bars = 5 µm.

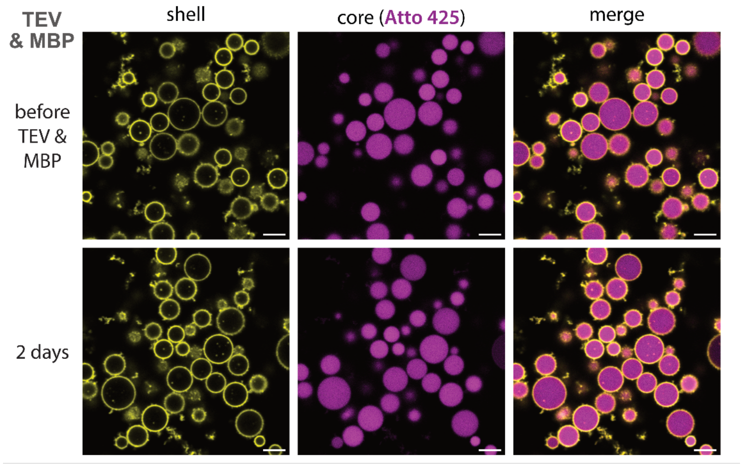

**Supplementary Fig. 3: Simultaneous addition of TEV protease and MBP does not induce PS.** Scale bars = 5 µm.

#### 4. MBP-FUS-GFP partitioning, FRAP and overview images in pristine PN

The partition coefficient P is defined as the ratio of GFP intensity in the PN core to that in solution. As shown in Supplementary Fig. 4a P is low for loading ratio, LR = 0.10 (P = 91), reaches its peak at LR = 0.30 (P = 448) and falls off continuously from there. The peak value for LR = 0.30 aligns well with the previously determined value in Figure 4d ($P_{pristine}$ = 410, LR = 0.30), with small variation expected due to PN batch-to-batch variations. The decline in P for higher LR is attributed to saturation behavior, which is more clearly visualized by plotting the product of LR and P, the enrichment-weighted loading (Supplementary Fig. 4b). While the relative P-value falls steadily, the absolute amount of protein in the PN core is still rising with higher LR. In case [FUS] reaches a maximum in the PN, the plot of LR x P would turn constant, which is not yet the case in our experimental window and explains the continuous change in PS morphologies with LR.

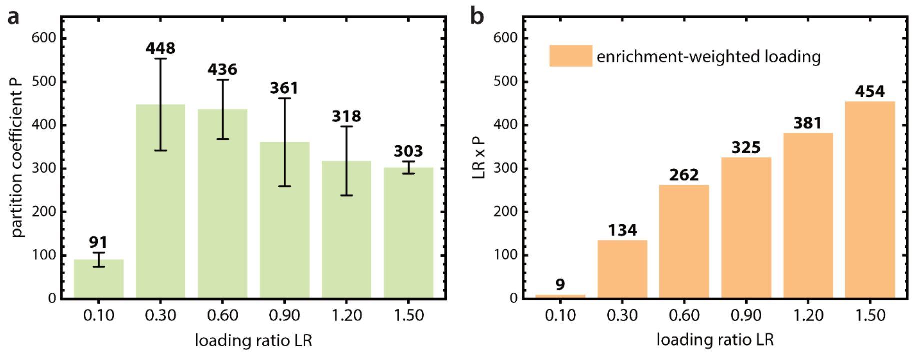

**Supplementary Fig. 4: Partitioning of MBP-FUS-GFP in pristine PN with different LR. a** Partition coefficients and their standard deviation as determined by overview CLSM images before MBP cleavage. N = 89, 87, 71, 32, 30, 18 (left to right). N denotes the number of individual PN compartments. **b** The product of LR and P plotted against LR yields the enrichment-weighted loading.

The dynamics of MBP-FUS-GFP inside pristine PN are dependent on the LR. Supplementary Fig. 5a shows the FRAP recovery curves of all LR shown in Figure 1. The general trend is a slower recovery with increasing LR, with the exception of LR = 0.90, which we view as outlier in this analysis. To quantify this behavior, we fitted the recovery curves and show the resulting half-recovery times $t_{1/2}$ and mobile fraction (endpoint recovery, ER) in Supplementary Fig. 5b. As expected, $t_{1/2}$ increases with LR and the mobile fraction remains stable between 40-50%. For details on FRAP analysis see Section 9 of the SI.

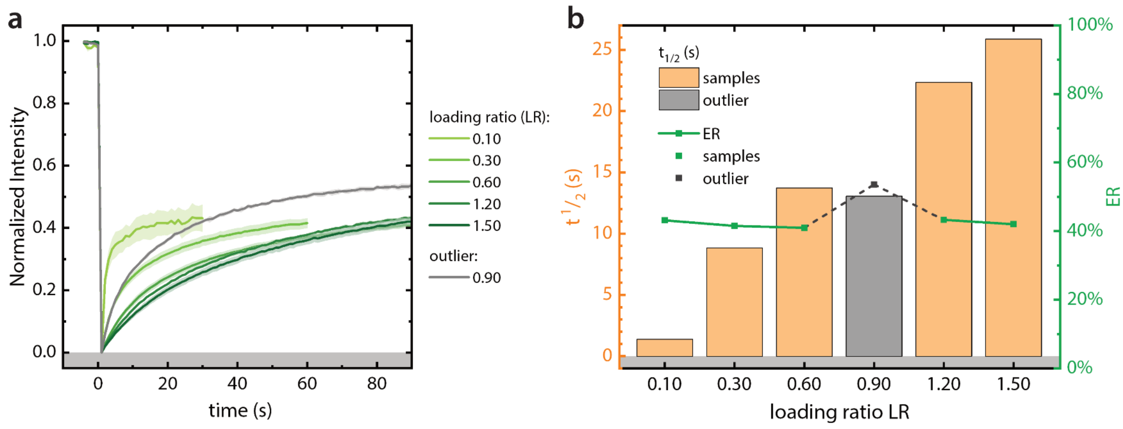

**Supplementary Fig. 5: FRAP analysis of pristine PN before MBP-cleavage. a** Recovery curves of MBP-FUS-GFP in pristine PN with different LRs. For each sample 3-4 recorded FRAP measurements were averaged into one curve (LR 0.10 - 0.60: N=4, LR 0.90 - 1.50: N=3, N denotes the number of individual PN compartments). The shaded area behind the curves represents the standard deviation. The recovery of LR=0.90 does not follow the general trend and has been marked an outlier. **b** Comparison of $t_{1/2}$ and endpoint recovery (ER) as resulting from fitting the average FRAP curves.

To provide an overview of the morphologies seen in all 6 selected LR samples after MBP cleavage, we provide the overview images corresponding to Figure 1 below the representative snapshot we have chosen.

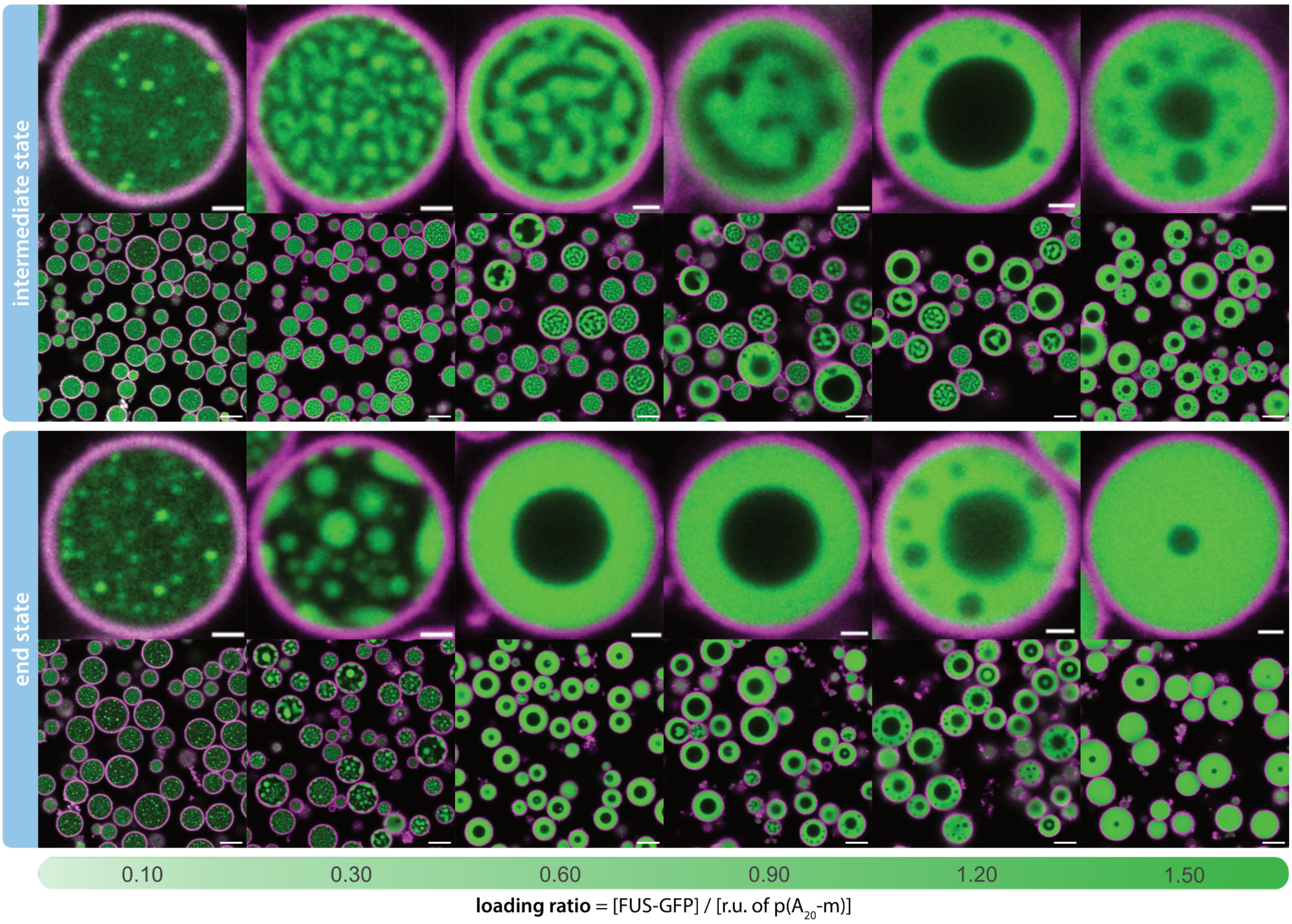

**Supplementary Fig. 6: Full view images of PS in pristine PN next to selected snapshots shown in Figure 1.** Scale bars: top rows (snapshots) = 1 µm; bottom rows (full view) = 5 µm.

#### 5. Numerical Simulations

The following describes the theoretical model and workflow for the numerical simulations underpinning Figures 2 and 3 of the main manuscript.

##### 5.1. Free energy

We write the free energy contributions in main text Equation 1 as:

$$F_{CH} = \sum_{k=1}^{n} \int dr \left[ A\varphi_k^2 (1-\varphi_k)^2 + \lambda_k^2 (\nabla \varphi_k)^2 \right] \tag{4}$$

$$F_{com} = \sum_{k=1}^{n} \left[ B_k \left( 1 - \frac{1}{\pi R_0^2} \int dr\, p(\varphi_k) \right)^2 \right] \tag{5}$$

$$F_{rep} = \frac{1}{2}\omega \sum_{k=1}^{n} \sum_{l=1}^{n} \left\{ (1-\delta_{kl}) \int dr \left( p(\varphi_k)\, p(\varphi_l) \right) \right\} \tag{6}$$

$$F_{mix} = \int dr \left[ f_{mix}(\{c_i\}) + \frac{1}{2} \sum_{i=1}^{m} \sum_{j=1}^{m} (\kappa_{ij} \nabla c_i \cdot \nabla c_j) \right] \tag{7}$$

These contributions represent an extended Cahn-Hilliard free energy ($F_{CH}$), PN compressibility ($F_{com}$), a penalty ($F_{rep}$) associated with spatial overlap between PN and a mixing free energy ($F_{mix}$), with

$$f_{mix}(\{c_i\}) = \sum_{i=1}^{m-1} \frac{c_i}{L_i} \ln c_i + \frac{\left(1 - \sum_{i=1}^{m-1} c_i\right)}{L_m} \ln \left( 1 - \sum_{i=1}^{m-1} c_i \right) + \frac{1}{2} \sum_{i=1}^{m} \sum_{j=1}^{m} (c_i c_j X_{ij}) \tag{8}$$

a multicomponent Flory-Huggins local mixing free energy with $L_i$ relative molecular size and

$$X_{ij}(\{\varphi_k\}) = \left( 1 - \sum_{k=1}^{n} p(\varphi_k) \right) \chi_{ij}^{(\alpha)} + \sum_{k=1}^{n} \left( p(\varphi_k)\, \chi_{k,ij}^{(\beta)} \right) \tag{9}$$

a spatially varying an interaction parameter which interpolates between extremes $\chi_{ij}^{(\alpha)}$ and $\chi_{k,ij}^{(\beta)}$ assigned to the regions outside and inside a PN. The solution is considered incompressible, so that $c_m = 1 - \sum_{i=1}^{m-1} c_i$. The sigmoidal function $p(\varphi_k)$ is defined as:[3,4]

$$p(\varphi_k) = \int_0^{\varphi_k} d\varphi_k' \left[ \varphi_k^2 (1-\varphi_k)^2 \right] / \int_0^1 d\varphi_k \left[ \varphi_k^2 (1-\varphi_k)^2 \right] \tag{10}$$

interpolates between the inside and outside of a PN and assures that in equilibrium $\varphi_k$ is zero/unity outside/inside PN indexed $k$, by obeying $d_{\varphi_k} p(\varphi_k) = d_{\varphi_k \varphi_k} p(\varphi_k) = 0$. The parameters $A$, $B$ and $\omega$ are dimensionless positive energy scales. Supplementary Equation (5) places a soft constraint to the size of a PN with $R_0$ representing a target radius. Lastly, $\lambda^2$ and $\kappa_{ij}$ are dimensionless coefficients penalizing gradients in, respectively, order parameters and concentrations. Gradient cross contributions are omitted for simplicity.

We introduce a novel approach to obtain a suitable evolution equation for the solute concentration. The main challenge is that for solute(s) and solvent to follow cell motion, the change in their local concentration, being a *conserved order* *parameter*, is subject to a velocity field $u=\sum_k u_k$ determined by changes in the *non-conserved* order parameters $\varphi_k$ (see main text Equation 3). As a result, $u$ is not necessarily non-divergent, implying that by bluntly coupling $c$ to $u$ through in a convection-diffusion equation, solute might spuriously form or disappear as the cells contract and extend, which they would do naturally. For this reason we decompose $u$ using a Helmholtz decomposition: $u_s=u-\nabla D$, with $u_s$ the non-divergent (solenoidal) and $\nabla D$ the curl-free component.

#### 5.2. Short range wall interaction

We crudely define a free energy density due to interaction of any of the components with the inner wall of the PN as:

$$F_{wall}=\int dr\left[\left(\chi_{DNA}^{(w)}c_{DNA}^{x}+\chi_{FUS}^{(w)}c_{FUS}^{x}+\chi_{solvent}^{(w)}c_{solvent}^{x}\right)p'(\varphi_0)\right] \tag{11}$$

, where $\varphi_0=1-\sum_{k=1}^{n}\varphi_k$ and $p'(\varphi_0)$ identifies the interfacial region. Supplementary Equation (11) does not represent a boundary condition, but rather represents a bulk contribution which only becomes important in regions where the gradient in the phase parameter is steep. The exponent $x$ tunes the concentration dependence and determines to what extent a component penetrates the interfacial region of the PN. Its value has no large effect on the PS dynamics. For all calculations we use $x$ = 0.25. Since the PN are rigid, we do not include the effect of this wall free energy in the evolution of the phase fields. We only set $\chi_{DNA}^{(w)}$ non-zero and implicitly allow for FUS accumulation at the wall through its attractive interaction with the DNA. Furthermore, we avoid large (negative) values to avoid pinning effects during the numerical simulations.

#### 5.3. Calculation details and input parameters

Numerical simulations involve numerically integrating dynamic Equations 2 and 3 (main text) in both the generation and the quenching stage on a square grid with periodic boundary conditions, using a forward-time-central-space (FTCS) explicit Euler finite difference scheme. All input parameters are dimensionless and listed in Supplementary Table 3 and Supplementary Table 4.

**Supplementary Table 3: Input parameters for numerical simulations.**

| | | |
|---|---|---|
| $N$ | 128 | domain size (grid length) |
| $m$ | 3 | number of components |
| $n$ | 3 | number of PN |
| $V_0$ | 0.65 | fractional target volume |
| $A$ | 1.0 | CH barrier |
| $B$ | 750 | PN compressibility |
| $\omega$ | 20 | PN overlap penalty |
| $\lambda^2$ | 1.5 | phase gradient energy |
| $\kappa_{ii}$ | 0.075 | concentration gradient energy |
| $\kappa_{ij}$ | 0 | concentration gradient energy |
| $L_{DNA}$ | 3 | relative size |
| $L_{MBPFUS}$ | 3 | relative size |
| $L_{FUS}$ | 3 | relative size |
| $L_{solvent}$ | 1 | relative size |
| $\chi_{DNA-MBPFUS}^{(\alpha)}=\chi_{DNA-MBPFUS}^{(\beta)}=\chi_{DNA-FUS}^{(\alpha)}=\chi_{DNA-FUS}^{(\beta)}$ | -2.75 | binary interaction |
| $\chi_{DNA-solvent}^{(\alpha)}$ | 4 | binary interaction |

| | | |
|---|---|---|
| $\chi^{(\beta)}_{DNA\text{-}solvent}$ | 0.3 | binary interaction |
| $\chi^{(\alpha)}_{MBPFUS\text{-}solvent} = \chi^{(\beta)}_{MBPFUS\text{-}solvent}$ | 0.8 | binary interaction |
| $\chi^{(\alpha)}_{FUS\text{-}solvent}$ | 1.2 | binary interaction |
| $\chi^{(\beta)}_{FUS\text{-}solvent}$ | 1.8 | binary interaction |
| $\chi^{(w)}_{DNA}$ | -0.15 | interaction inner wall |
| $\chi^{(w)}_{FUS}$ | 0 | interaction inner wall |
| $\chi^{(w)}_{solvent}$ | 0 | interaction inner wall |
| $D_{DNA} = D_{FUS} = D_{solvent}$ | 8 | Diffusivity |
| $\xi$ | 0.125 | friction coefficient |
| $\Gamma$ | 1.0 | kinetic coefficient |
| $\Delta t$ | $2.5\times10^{-3}$ | integration time step |

Some remarks: i) the target radius of a PN (see Supplementary Equation (5)) follows from the total fractional target volume of PN material according to: $R_0 = V_0 \sqrt{N^2/m\pi}$, ii) we assign the size of both "solutes" three times that of a "solvent" particle as a compromise between retaining some asymmetry in the mixing free energy expected for polymer solutions and guaranteeing numerical stability at reasonably large time steps in our explicit integration scheme, iii) the following reduced concentration gradient energy matrix is obtained by applying the incompressibility condition: $\bar{\kappa} = \begin{pmatrix} 0.15 & 0.075 \\ 0.075 & 0.15 \end{pmatrix}$, and iv) since time scales are not absolute, for simplicity we assume the same self-diffusivity for all three components. The diffusion mobilities are derived from the molecular sizes and self-diffusivities according to fast-mode theory[5,6], as implemented in previous work.[7,8]

**Supplementary Table 4: Mean volume fractions per loading ratio (independent components).**

| | |
|---|---|
| LR = 0.37 | $\bar{c}^{(0)}_{DNA}$ = 0.1 |
| | $\bar{c}^{(0)}_{MBPFUS}$ = 0.04375 |
| LR = 0.43 | $\bar{c}^{(0)}_{DNA}$ = 0.1 |
| | $\bar{c}^{(0)}_{MBPFUS}$ = 0.05 |
| LR = 0.65 | $\bar{c}^{(0)}_{DNA}$ = 0.1 |
| | $\bar{c}^{(0)}_{MBPFUS}$ = 0.075 |
| LR = 0.80 | $\bar{c}^{(0)}_{DNA}$ = 0.1 |
| | $\bar{c}^{(0)}_{MBPFUS}$ = 0.1 |
| LR = 1.0 | $\bar{c}^{(0)}_{DNA}$ = 0.1 |
| | $\bar{c}^{(0)}_{MBPFUS}$ = 0.125 |
| LR = 1.3 | $\bar{c}^{(0)}_{DNA}$ = 0.1 |
| | $\bar{c}^{(0)}_{MBPFUS}$ = 0.165 |

#### 5.4. Discussion

The model predicts that upon cleaving off the MBP tag, an influx of FUS takes place from the surrounding medium into the PN, as the system lowers the free energy by driving FUS into regions with a low solvent fraction. Although this may happen in the experiments too, the model exaggerates the effect since the calculations avoid very small concentrations to guarantee numerical stability. To counteract the excessive influx, we set the FUS-solvent interaction parameter in the exterior region slightly lower than in the interior region (see Supplementary Table 3). The result nicely reproduces what is seen in the experiments, namely that the surface enrichment of the dense phase is more prominent for high LR compared to low LR.

We point to some unresolved differences between the computed and experimental morphologies at late stages. Especially at high loading ratios, the experiments show a ‘collapse’ of the dense phase network towards the inner wall of the PN. This is not seen in the computations. We suspect that the tethering of part of the DNA to the PN not only causes a strong drive for wetting of the inner wall, but possibly also gives rise to a long-range elastic field. In contrast, in its current form, the model only includes a short-range wall potential for the DNA (see SI Section 5.2) but does not include a real ‘wetting condition’, wherein this surface potential balances the penalty associated with the development of perpendicular concentration gradients, as we have implemented earlier[8]. In future work, we look forward to implement such refinements.

#### 6. Melting temperatures of core modification strands

The following Supplementary Fig. 7 shows the calculated melting curves for all core modification strands used in Figure 4. They have been calculated with Nupack 4.0.2.0 for the experimental concentrations and reveal melting temperatures, $T_m$, of above 62.1°C. Even for the modifier strand with the lowest melting temperature (**6**), the fraction of duplex at the experimental temperature (25 °C) is 100%, with more than 20 °C distance to the onset of melting. Since the modification strands are used sub-stoichiometrically (80% to target), the probability of unbound core modification strands in solution is minimal.

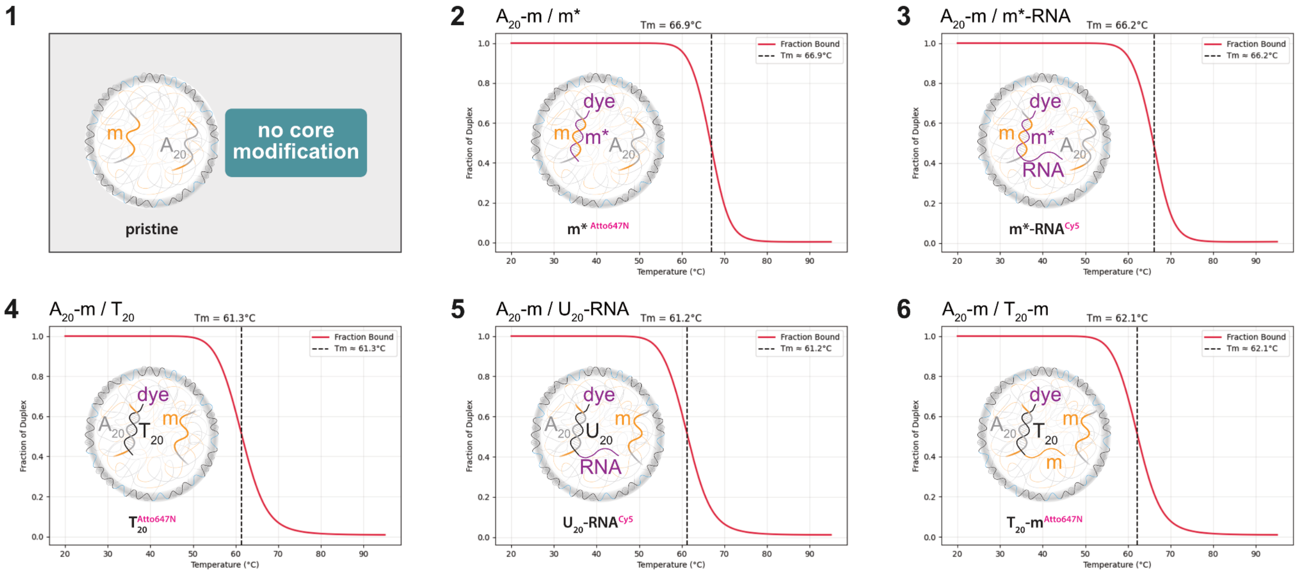

**Supplementary Fig. 7: Calculated melting curves for all core modification strands with the core repeating unit in experimental buffer.** Melting curves have been calculated with Nupack 4.0.2.0 at the following concentrations: [$A_{20}$-m] = 4.92 µM, [core mod.] = 3.94 µM, [$Na^+$] = 200 mM and [$Mg^{2+}$] = 10 mM.

#### 7. Overview images of typical PN sample to underscore homogeneity

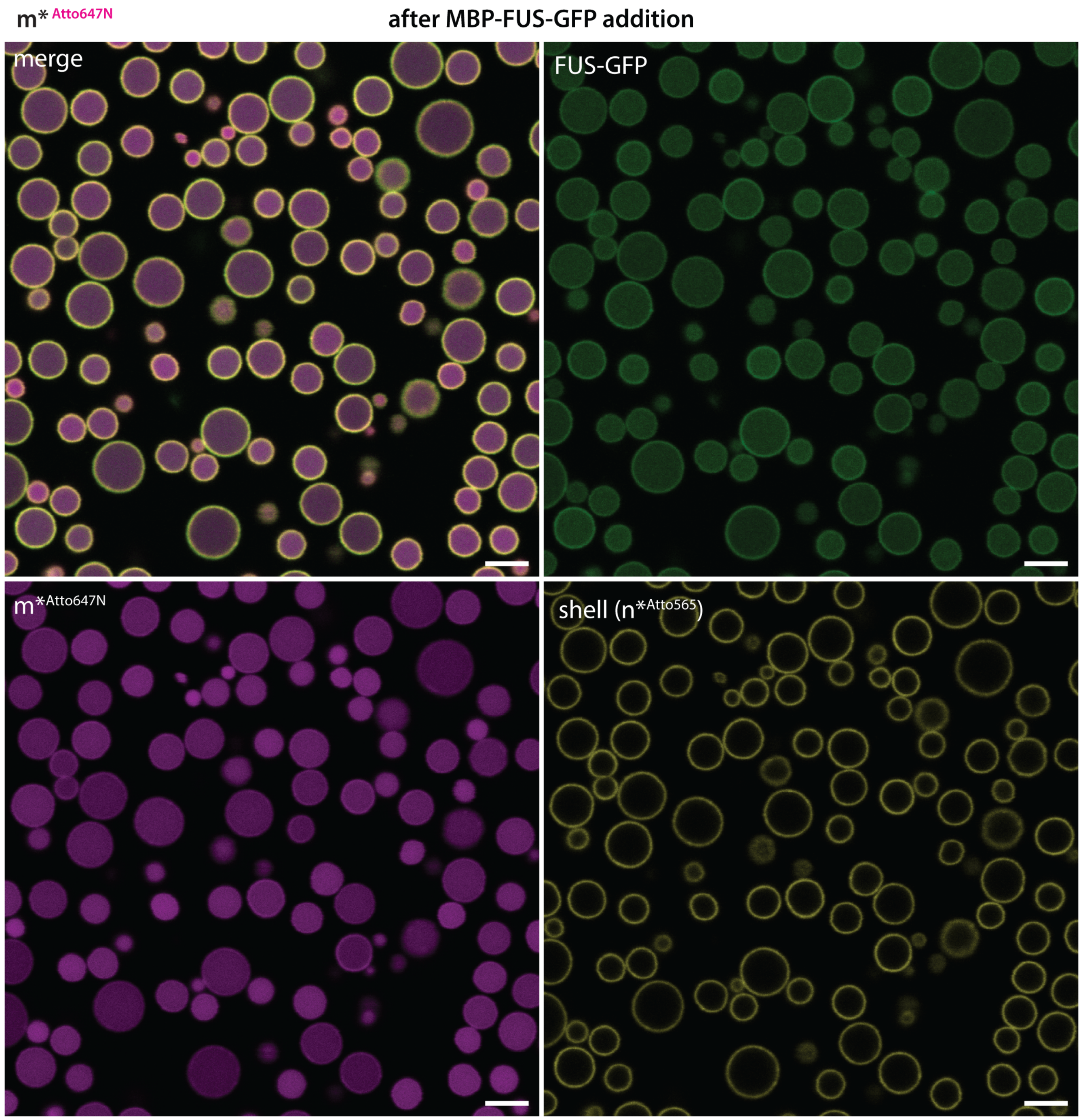

**Supplementary Fig. 8: Typical PN sample.** Representative CLSM image highlighting the low polydispersity of PN samples. Scale bars = 5 µm.

#### 8. Single channel CLSM images of sequence dependent PS and overview images

For increased visibility of the FUS and pA core channel, images show both isolated fluorophore channels. The shell channel is omitted.

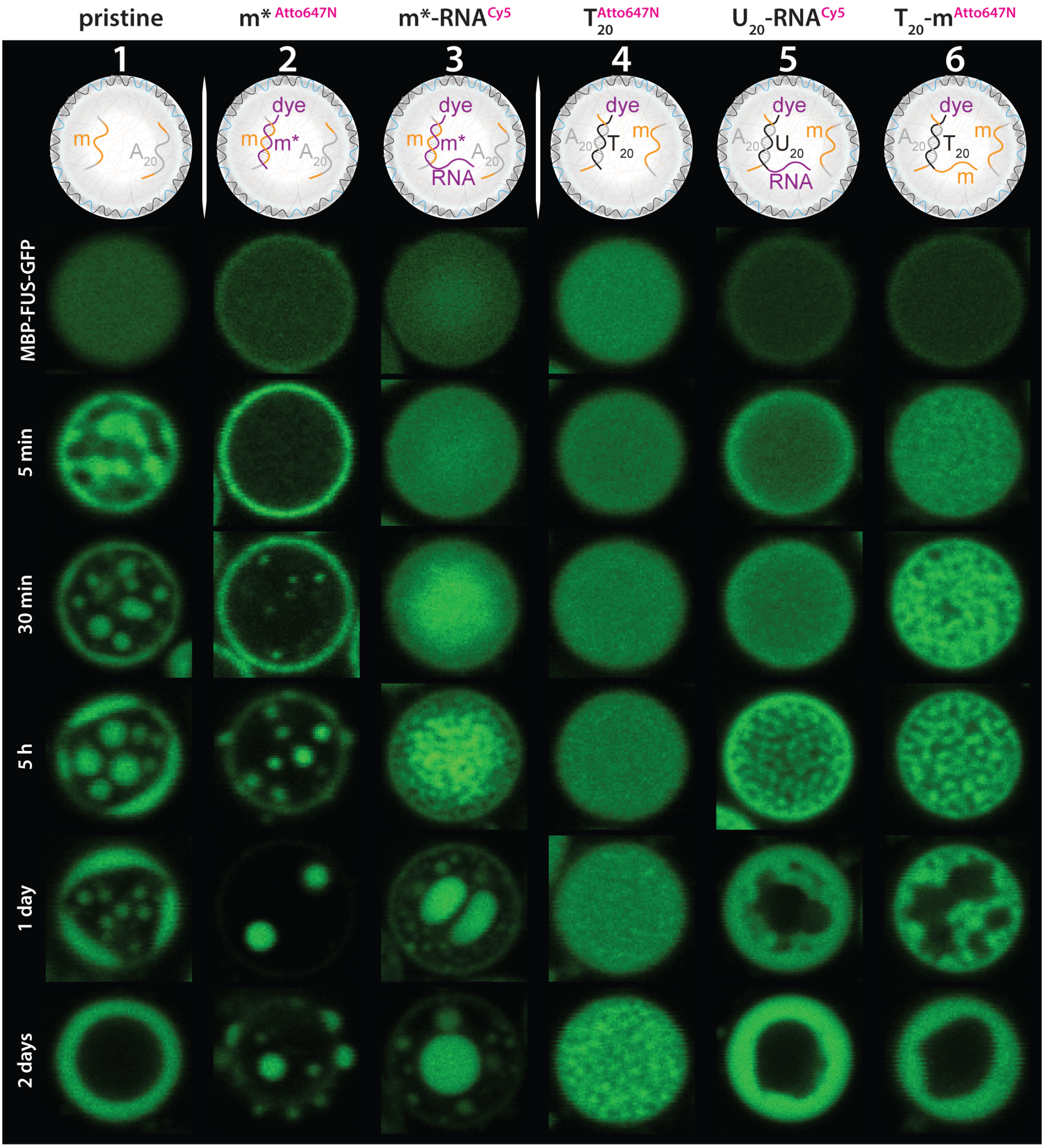

**Supplementary Fig. 9: Green channel for images shown in Figure 4c, which shows the fluorescent signal of the GFP-tagged FUS protein.**

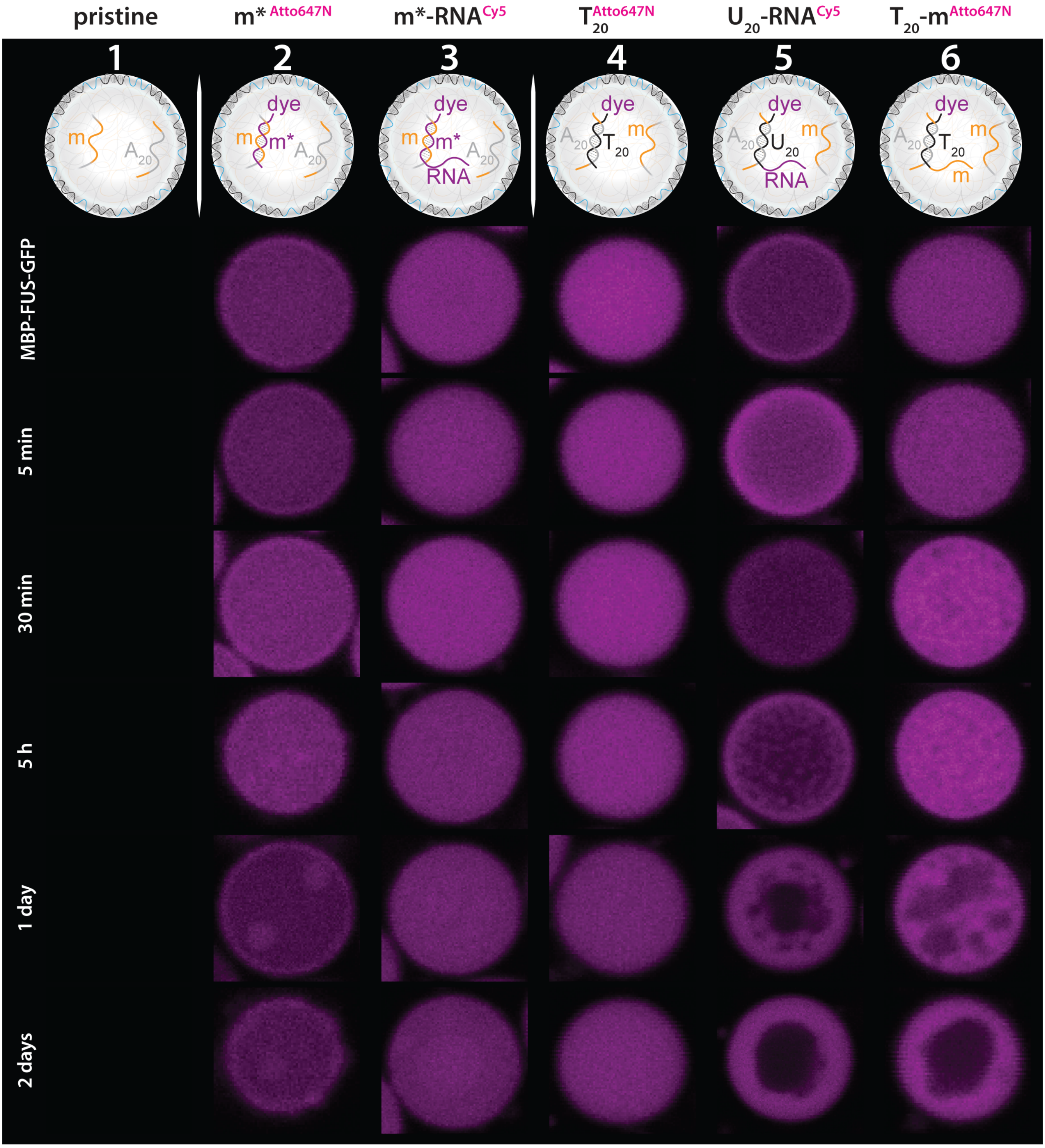

**Supplementary Fig. 10: Magenta channel for images shown in Figure 4c.** In all experiments but the first one we have added nucleic acid strands with a fluorescent dye, which is Cy5 for RNA and Atto 647N for DNA. They are hybridized with the PN core. Images show if they are drawn into the FUS-GFP condensates or not.

The following two figures show overview images for the representative PN that have been isolated in Figure 4c, marking the position of those PN with white boxes.

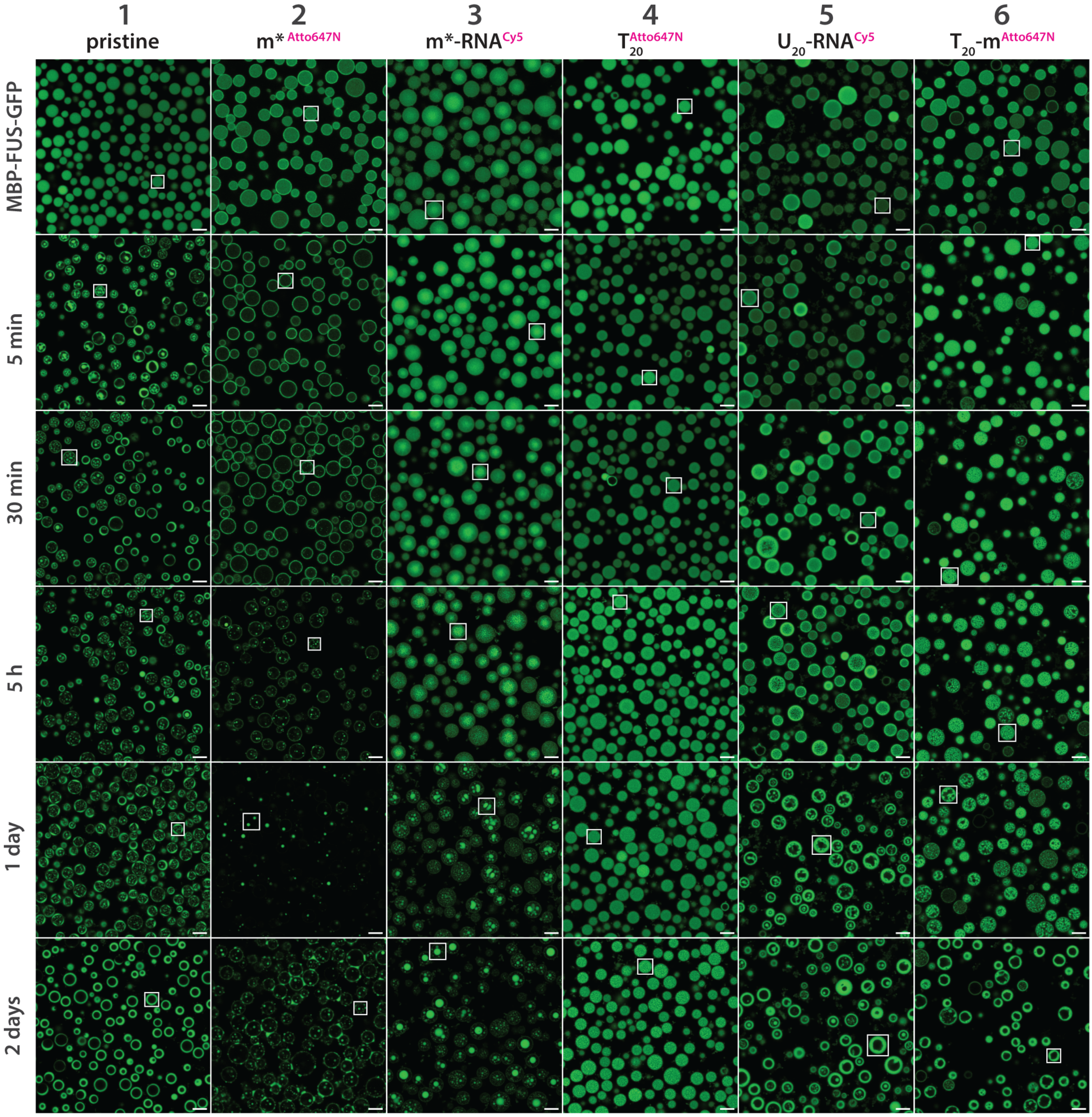

**Supplementary Fig. 11: Overview images for single PN shown in Figure 4c, with isolation of the green FUS-GFP channel.** The white boxes indicate the position of the PN chosen for Figure 4c. Scale bars = 5 µm.

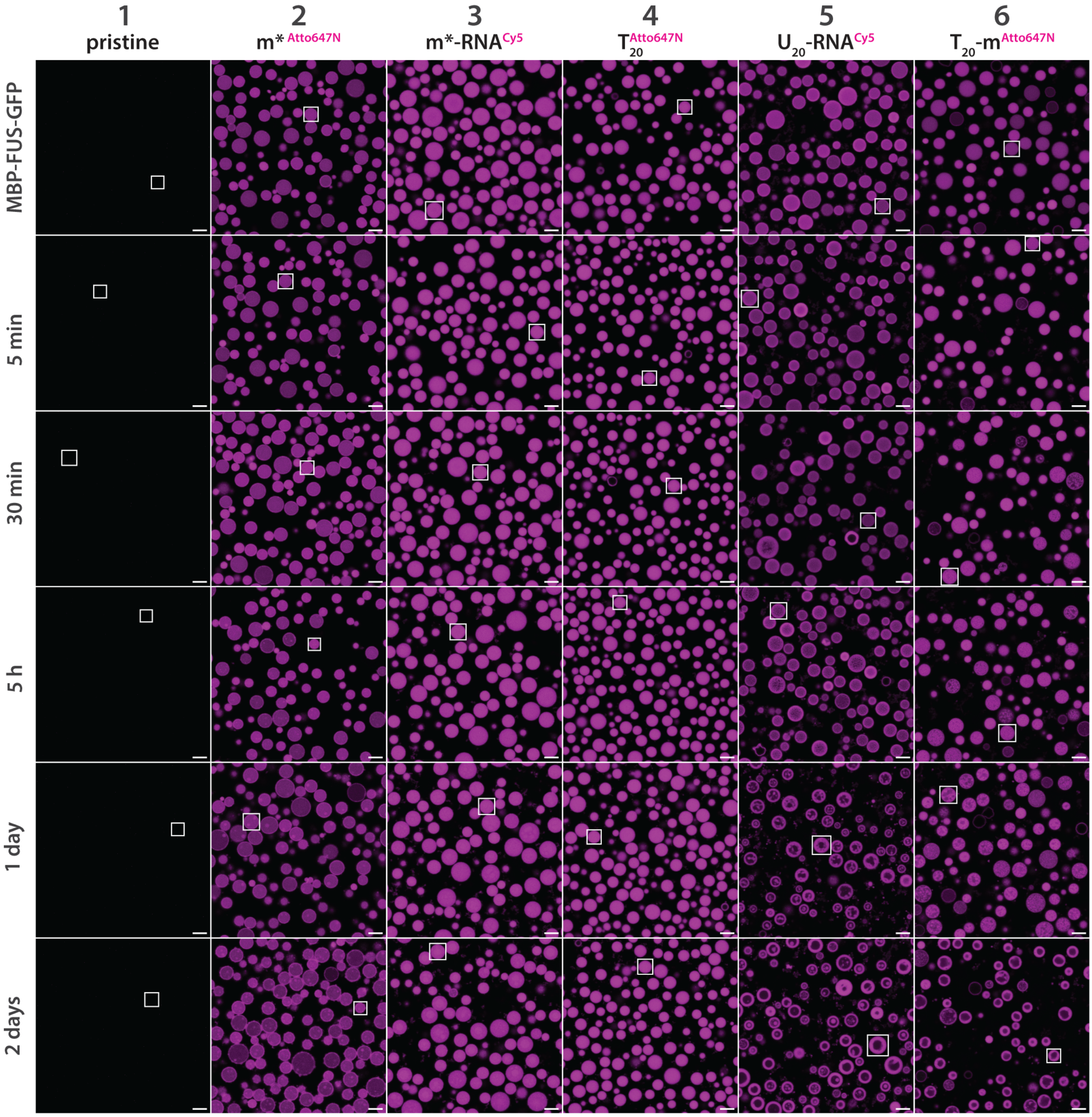

**Supplementary Fig. 12: Magenta channel of overview images for Figure 4c, showing the fluorescence of the core modification sequences.** Selected PN in Figure 4c are indicated with white boxes. Scale bars = 5 µm.

#### 9. Fluorescence recovery after photobleaching (FRAP)

##### Instrumentation, bleach setup & measurement

All FRAP experiments were conducted on a Leica Stellaris 5 CLSM. The bleaching procedure involved the selection of an approximately 1-2 µm sized ROI in the center of a PN, which was bleached with the 488 nm laser at 100% intensity for 0.42 s. The recovery was followed in time intervals of 1 s per frame, which is a compromise of high sampling rate and avoidance of sample bleaching.

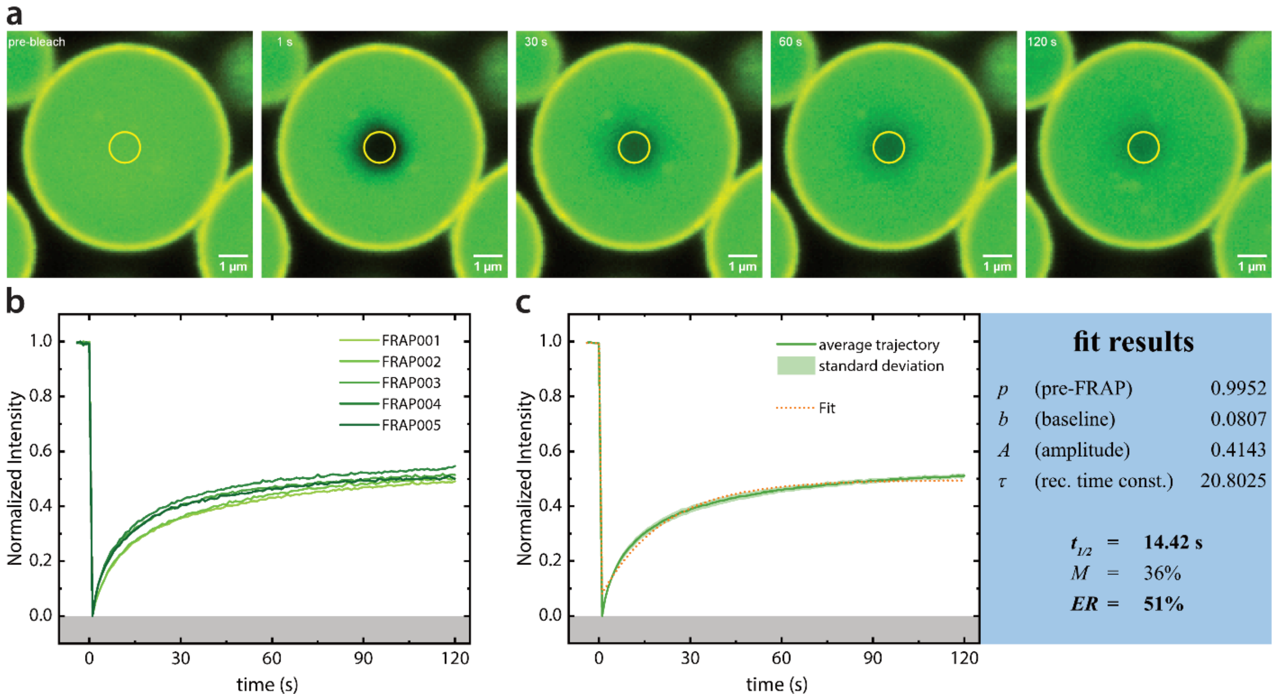

**Supplementary Fig. 13: Bleach setup, time progression and analysis. a** FRAP measurement of a representative sample, 75% XL before TEV addition. One bleach ROI per PN is aligned to the center of the PN as a partial bleach of the FUS-GFP channel. The yellow circle shows the manually chosen ROI for analysis of the intensity before and after bleaching, which is not congruent with the actual bleach region. **b** Combined plot of five FRAP traces with a Min-Max normalization of the intensity. The sample shown in **a** is plotted as FRAP001. **c** Average trajectory of the combined plots, shaded with the standard deviation. Orange dotted line shows the exponential fit function (Supplementary Equation (12)) for the recovery curve, which yields the fit results shown on the right.

##### FRAP analysis and curve fitting

The data was processed and fitted with the Stowers FRAP plugin, using the batch FRAP fit jru v1 function. For intensity measurements only raw images are used. The plugin uses the following fitting function, which is the standard exponential FRAP fit.

$$I(t) = b + A\left(1 - e^{-\frac{t}{\tau}}\right) \tag{12}$$

From this we obtain the baseline intensity *b*, the amplitude of the recovery curve *A* and the recovery time constant *τ*. From the fit results we obtain $t_{1/2}$ and the mobile fraction *M* with the following calculations, which also involve the average pre-FRAP intensity *p*.

$$t_{1/2} = \tau \times \ln(2) \tag{13}$$

$$M = \frac{A - b}{p - b} \tag{14}$$

However, several FRAP traces do not reach a reliable plateau within the imaging window. In these cases, the recovery extent is quantified directly from the normalized fluorescence trace at the final recorded time point rather than from fit-derived mobile fractions. We report this value as the endpoint recovery *ER*.

#### 10. FRAP in PN with modified core

The following Supplementary Fig. 14 shows the FRAP images corresponding to Figure 4f. As reflected in the intensity traces in the main text, m*$^{Atto647N}$ is the only sample showing recovery in the first 20 seconds of observation.

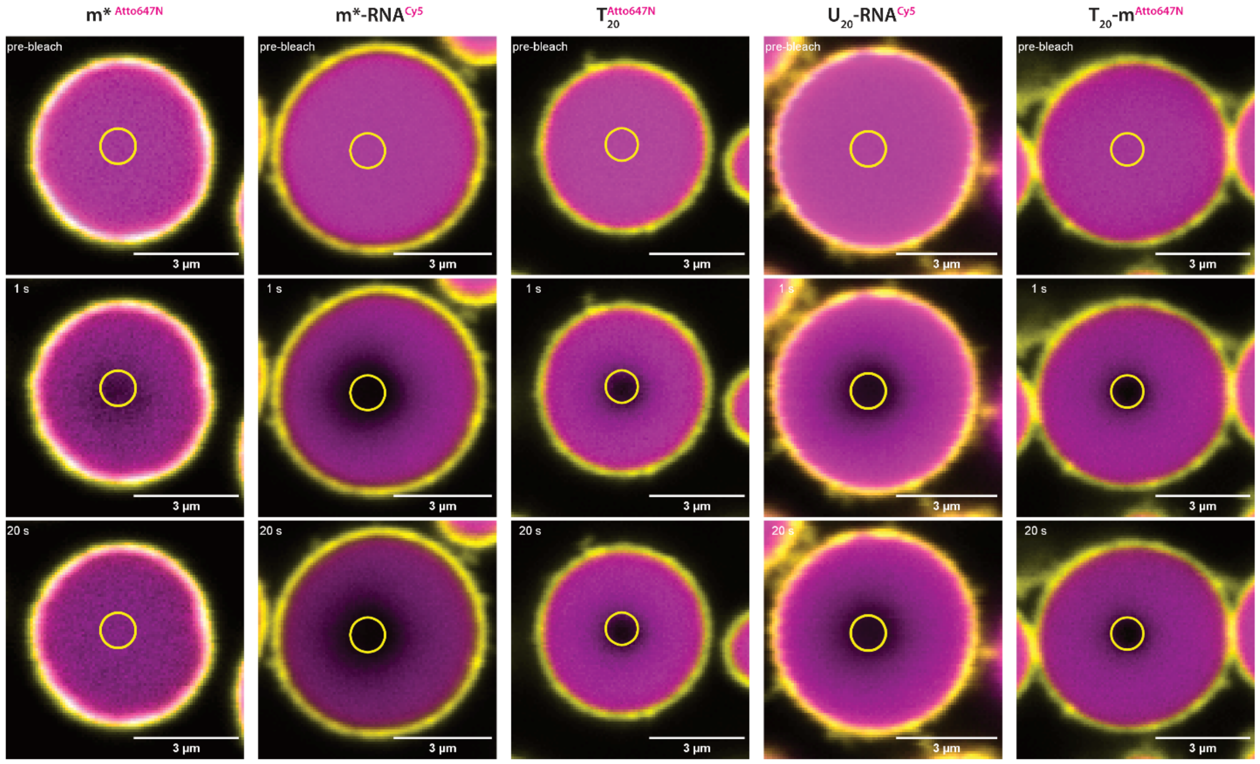

**Supplementary Fig. 14: PN before bleaching, immediately afterwards and recovery progression for core modified PN in Figure 4f.** All images are adjusted in contrast with the same boundaries. The small yellow circle shows the region of interest (ROI) used for the intensity measurements, which are always performed with raw images.

#### 11. All-atom MD simulations of $T_{20}$ and $A_{20}$ binding to RGG2

FUS RGG2 is arginine-rich and engages nucleic acids through a combination of transient and sequence-dependent interactions. Here, we compare complexes with $A_{20}$ and $T_{20}$ DNA using a continuous 1 μs trajectory and nine independent 100 ns replicates to capture both long-timescale behavior and seed-to-seed variability. We note that short replicate simulations are mostly useful to report on local contacts and are likely too short to sample the overall conformational dynamics of $A_{20}$ and $T_{20}$. Short simulation duration can severely bias the calculations on structural observables, in absence of kinetic models that correct for such biases.[9] To determine whether $A_{20}$ and $T_{20}$ differ in overall stability and in their relative protein–DNA arrangement, we first analyzed the end-to-end distance of the ssDNA.

##### 11.1. Conformational stability

To compare the global conformational behavior of FUS RGG2 in complex with $T_{20}$ and $A_{20}$, we analyzed the end-to-end distance of the ssDNA. The end-to-end distance was defined as the distance between the oxygen atom of the 5′ hydroxyl group and the oxygen atom of the 3′ hydroxyl group. This metric directly quantifies the overall extension of the DNA chain and thereby reports on its base-stacking propensity and compactness.

As shown in Supplementary Fig. 15a, $A_{20}$ adopts a markedly more extended conformation than $T_{20}$ throughout the 1 μs continuous simulation. The higher end-to-end distance of $A_{20}$ is fully consistent with its stronger intrinsic base-stacking tendency, which favors a more rigid, extended single-stranded structure. In contrast, $T_{20}$ forms more compact conformations on average, reflecting its lower stacking propensity and greater flexibility. Note, that the end-to-end distance of the replicate simulation did not show such a clear difference, with average end-to-end distance of 46.26 Å (standard error of the mean = 0.99 Å) for $A_{20}$ and 44.23 Å (standard error of the mean = 1.98 Å) for $T_{20}$. We note that Pollack et al have observed, by combining SAXS experiments and atomistic modeling, that overall extension of poly(A) and poly(T) differs only a little, while poly(A) has higher stacking propensity than poly(T).[10]

Representative snapshots from the continuous trajectories (Supplementary Fig. 15b,c) illustrate the structural consequences for the protein–DNA arrangement: $T_{20}$ remains wrapped more closely around the protein surface, whereas $A_{20}$ displays DNA segments that extend farther away from the protein. Taken together, these data indicate that $T_{20}$ forms a more compact and consistent FUS–DNA arrangement, while $A_{20}$ exhibits a more extended and dynamic DNA conformation. This behavior aligns with previous atomistic simulations of poly(A) and poly(T) ssDNA, which showed the same qualitative difference in chain extension driven by base stacking.[10–12] The increased extension and self-stacking of $A_{20}$ reduce base accessibility for transient protein contacts, providing a structural rationale for the sparser and more variable interaction network observed with $A_{20}$ compared with the broader, more coherent interface formed with $T_{20}$.

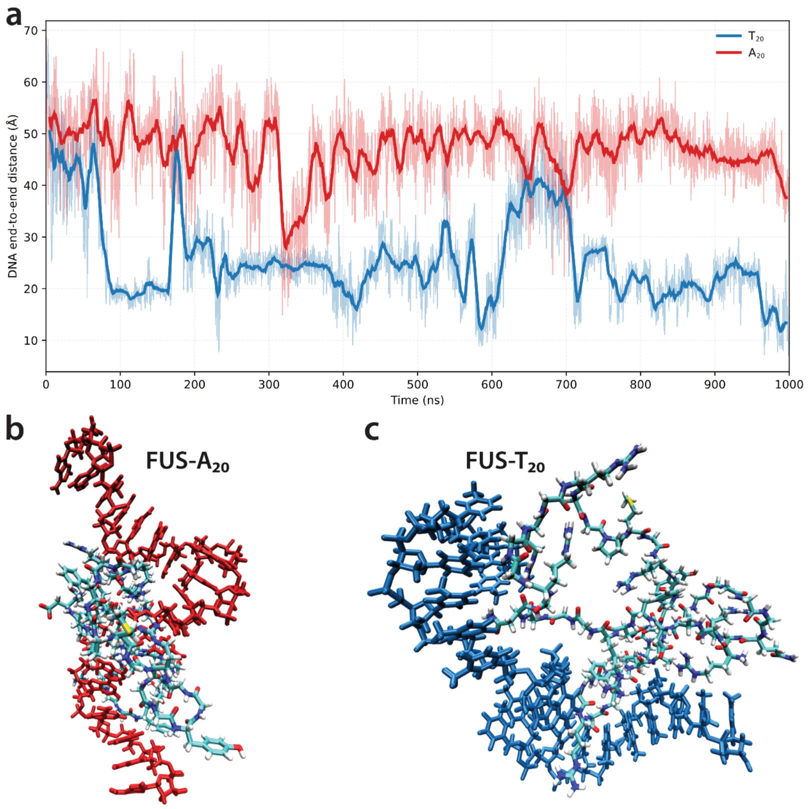

**Supplementary Fig. 15: Conformational stability and DNA positioning in FUS-DNA simulations. a** End-to-end distance of the DNA in complexes with $A_{20}$ (red) and $T_{20}$ (blue) during the 1-µs continuous simulations. Thick lines represent a running mean and faint lines the raw trajectory. **b,c** Representative snapshots from the continuous trajectories showing the spatial arrangement of DNA relative to FUS for the $A_{20}$ (red) and $T_{20}$ (blue) complexes. Structures were visualized in VMD.

#### 11.2. Cation-π interactions

Given that FUS RGG2 is especially rich in arginine, cation-π interactions are of particular interest. As shown in the continuous trajectory contact maps (Supplementary Fig. 16a), $T_{20}$ forms substantially stronger and more focused cation-π hotspots than $A_{20}$ (max pair occupancy 65% vs 18%). $T_{20}$ also forms many cation-π interactions in the replica simulations. This supports the idea that cation-π interactions as seen in the 1 µs trajectory are relevant for the interactions of disordered regions of FUS with poly(T). On the level of replicate simulations across independent seeds, many cation-π interactions are also observed for $A_{20}$. We note that strongest replicate hotspots in $A_{20}$ are almost double of those observed for $T_{20}$ (10% vs 5%, Supplementary Fig. 16b), yet, short simulation durations make it difficult to be certain about these differences. Both systems display large seed-to-seed variability, consistent with the transient nature of these contacts. To give a structural example of why cation-π can be very trajectory-dependent, a representative snapshot in Supplementary Fig. 16c-d shows $A_{20}$ containing multiple stacking-like stretches that can reduce base-ring accessibility for arginine, while $T_{20}$ shows less stacking and remains more exposed. Overall, the cation-π contacts show a trajectory-dependent contrast. $T_{20}$ can form highly persistent hotspots in the long run, whereas $A_{20}$ shows higher average occupancy across replicate sampling.

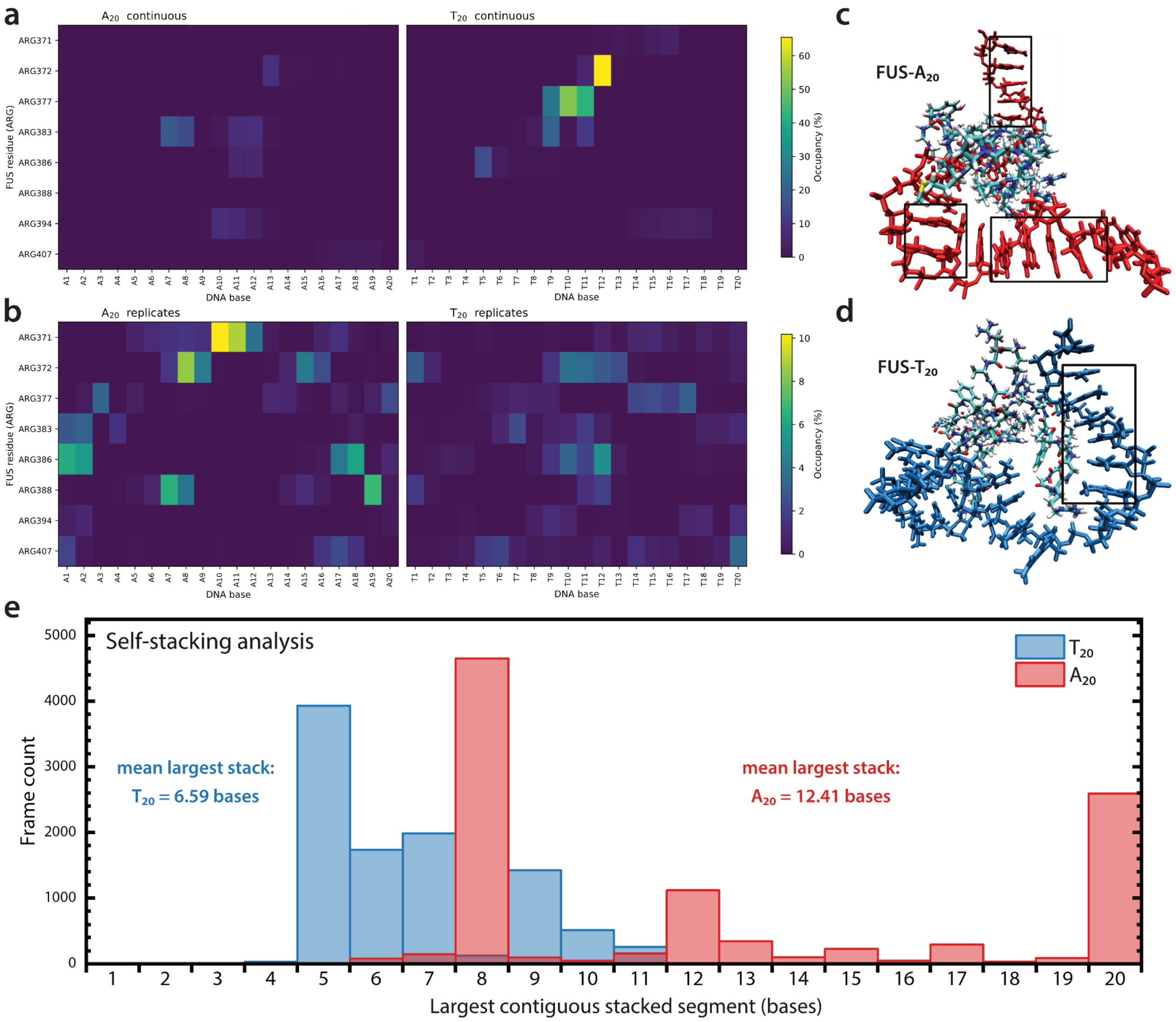

**Supplementary Fig. 16: Cation-π interaction maps and representative base presentation in FUS-DNA complexes. a** Heatmaps show the occupancy of cation-π contacts between FUS arginine residues and DNA bases for the continuous 1-μs simulations and **b** as the mean across nine independent 100-ns replicate simulations of the $A_{20}$ and $T_{20}$ systems. Shared color scales within each row allow direct comparison between the two DNA sequences. **c-d** Representative snapshots from the continuous trajectories of FUS RGG2 bound to $A_{20}$ (**c**, red) and $T_{20}$ (**d**, blue). Boxed regions highlight base arrangements with a stacking-like presentation that limit exposure of aromatic base surfaces for arginine cation-π contacts. Structures were visualized in VMD. **e** Distribution of the largest contiguous stacked segment per frame shows that $A_{20}$ forms substantially longer self-stacked stretches than $T_{20}$, consistent with reduced base accessibility for FUS binding.

Adenine being larger and having a larger π-system could suggest stronger binding at first glance, but the results support a different balance. The key is that increased base surface does not necessarily translate to increased protein interface if the bases prefer to interact with each other. Stronger adenine self-stacking can shift surface area into base–base interactions and reduce base accessibility for the distributed, transient contacts that an arginine/glycine-rich disordered domain typically uses. This is different in poly(T), because it is less prone to self-stacking, meaning the bases stay more exposed and are easier to access for protein contacts. This supports a broader multivalent interface, stronger water-mediated connectivity, and in some trajectories very persistent hotspots.

To quantify this effect, we additionally analyzed contiguous intra-strand base stacking along the $A_{20}$ and $T_{20}$ oligomers over the MD trajectories (Supplementary Fig. 16e). This analysis reveals that $A_{20}$ populates substantially longer self-stacked segments than $T_{20}$, consistent with reduced nucleobase accessibility for interaction with FUS. The distribution of the largest contiguous stacked segment per frame was strongly shifted toward larger values for $A_{20}$ (mean 12.41 bases) compared with $T_{20}$ (mean 6.59 bases), supporting the interpretation that adenine self-stacking limits protein-accessible base surface. Additionally, stacks of at least 10 bases occur in 50.3% of $A_{20}$ frames, but only 7.8% of $T_{20}$ frames. In the short replica trajectories this trend was not seen, with median number of stacked bases 5.7 for $A_{20}$ and 6.3 for $T_{20}$. This is also in line with the end-to-end distances of ~46.2 Å and ~44.2 Å for $A_{20}$ and $T_{20}$. We speculate that the limited simulation duration of the replicate simulations means that its extension and stacking cannot be accurately determined from these short simulations.

Previously, the stronger stacking of poly(A) as compared to poly(T) has been linked to the stronger binding of T to single-stranded DNA binding proteins.[13] Such stronger stacking has been revealed by combining SAXS experiments and atomistic modeling[10,11] and extensive atomistic molecular dynamics simulations of single-stranded DNA in absence of proteins, but also in atomistic modeling of poly(A) and poly(U) RNA.[12] Our simulations suggest these differences could also be relevant for the interactions of disordered proteins with T and A in DNA and the stronger ability of T to interact with FUS.

#### 11.3. Simulation setup

##### Input Sequences and Model Generation

To compare sequence-dependent interactions of FUS (UniProt P35637) RGG2 with ssDNA, the RGG2 segment (residues 371-421) was simulated in complex with two 20 nucleotide long ssDNA homopolymers: $A_{20}$ and $T_{20}$. The initial structure for RGG2 was generated with Hierarchical Chain Growth (HCG)[12,14–17] with default parameters and 50 output models, with the initial protein conformation being selected from this ensemble. The ssDNA 20-mers were generated using DNA Sequence to Structure (SCFBio IITD) with B-DNA settings[18]. The selected RGG2 model was combined with either $A_{20}$ or $T_{20}$ by combining the two structures into a single PDB file, without performing additional docking.

##### MD Simulation Workflow (GROMACS)

All-atom MD simulations were performed to sample the dynamics of the RGG2-ssDNA complexes and enable time-resolved interaction analysis. All systems were prepared in GROMACS 2024.4 using pdb2gmx to assign force-field parameters and add missing hydrogens. The simulations used the the amber14sb_OL15 force field with the TIP4P water model, as amber14sb_OL15 is a widely used parameter set for protein-nucleic acid simulations, and TIP4P as the associated explicit solvent model. Each complex was placed in a dodecahedral simulation box with a minimum distance of ≥ 1.0 nm between any protein or DNA atom and the box edge. Each complex was solvated with TIP4P water molecules and $Na^+$ and $Cl^-$ ions were added to neutralize the system and reach 0.15 M NaCl. Exact box dimensions and composition for each complex are listed in Supplementary Table 5. Both systems were relaxed by steepest-descent energy minimization to remove steric clashes. The systems were each equilibrated in two steps: first, NVT was run with 100 ps at 300 K with position restraints on the solute, using the V-rescale thermostat and fixed box volume to equilibrate temperature. Second, NPT was run with 100 ps at 300 K and 1 bar with position restraints maintained, using V-rescale temperature coupling and an isotropic Parrinello-Rahman barostat to relax box volume and solvent density. The production simulations were run under NPT conditions at 300 K and 1 bar without positional restraints. All runs used the leap-frog integrator with a 2 fs time step and LINCS constraints on bonds to hydrogen. Long-range electrostatics were treated with particle-mesh Ewald (PME), Lennard-Jones interactions used cutoffs consistent with the selected force field and periodic boundary conditions were applied in all directions. For each system (RGG2-$A_{20}$ and RGG2-$T_{20}$), one 1 μs trajectory per system was generated as 10 sequential chunks each 100 ns long. For the first chunk, initial velocities were assigned randomly in GROMACS (gen_seed = -1), and subsequent chunks were continued from the previous segment. In addition, an ensemble of nine independent replicas was simulated for 100 ns each using defined (non-random) velocity seeds 1001-1009 to standardize initial conditions between the two systems. Coordinates and energies/log files were written every 10 ps for downstream analysis.

**Supplementary Table 5: System composition and simulation box dimensions for the solvated RGG2-ssDNA complexes.**

| System | Box (nm) | Volume ($nm^3$) | Total atoms | Waters (SOL) | $Na^+$ | $Cl^-$ |
|---|---|---|---|---|---|---|
| RGG2-$A_{20}$ | 10.206 × 10.206 × 7.217 | ~751.65 | 97,837 | 24,110 | 80 | 68 |
| RGG2-$T_{20}$ | | | 97,913 | 24,129 | | |

##### Force Fields and Water Models

Both systems, FUS-$T_{20}$ and FUS-$A_{20}$, were simulated with the amber14sb_OL15 force field, which combines the AMBER ff14SB protein parameters with the DNA refinements of OL15.[19–21] ff14SB was developed to improve protein side-chain and backbone torsions relative to earlier AMBER protein force fields,[20] whereas OL15 is a modern DNA parameter refinement.[19,22] Simulations of all systems were performed with the original TIP4P water model.[21]

#### 11.4. Postprocessing, Analyses and Visualization

**Centering**

Artifacts from periodic boundary conditions (PBC) were corrected by first clustering the Protein-DNA complex (-pbc cluster) and then recentring the complex (-pbc mol -center -ur compact) using GROMACS trjconv prior to analysis.

**Analyses**

Analyses were run in Python with MDAnalysis[23,24] on PBC-corrected trajectories on the Mogon2 HPC cluster (CPU nodes): protein-DNA contacts, water-mediated bridging, RMSD metrics, and arginine-specific interaction geometries using fixed distance cutoffs.

**Cation-π proximity map**

To quantify arginine-base cation-π proximity, arginine-nucleotide pair occupancy maps were computed using a distance-based criterion. For each frame, an Arg-base pair was counted if the arginine side-chain cationic group was within 6.0 Å of the nucleobase aromatic ring (cut off = 6.0 Å). Each unique residue pair was counted at most once per frame, and occupancies were calculated as the percentage of analyzed frames (trajectory stride = 5 frames).

**Self-stacking analysis**

To quantify contiguous nucleobase stacking in the DNA strands, a cut-off based stacking analysis was performed on the PBC-corrected trajectories using Python with MDAnalysis. For each frame, stacking was evaluated between neighboring nucleobases along the strand (residues 1-2, 2-3, …, 19-20) using a distance-based criterion. A neighboring base pair was classified as stacked when the distance between the centers of mass of the nucleobase ring atoms was ≤ 4.0 Å under periodic boundary conditions. For adenine, the ring atoms N1, C2, N3, C4, C5, C6, N7, C8, and N9 were included; for thymine, the atoms N1, C2, N3, C4, C5, and C6 were used. Analyses were carried out with a stride of 10 frames.

**Software**

Simulations and trajectory preprocessing were performed with GROMACS 2024.4. All analyses were run on Mogon using a micromamba-managed Python environment (MDAnalysis + NumPy/Pandas) to generate the raw output files. The subsequent post-processing and figure generation were performed locally on MacBook using Python 3.11. For visualization all graphs were generated using MDAnalysis scripts, protein-DNA structure snapshots were created in VMD[25–32] and edited in Inkscape (version 1.4.2). Analysis and workflow scripts were prepared with assistance from ChatGPT and executed after manual verification.

All scripts and general workflow can be found on:

https://gitlab.com/antonia-18081/fus-ssdna/-/tree/d1403e61005a55e6097156b755f5dd57202846eb/.

#### 12. MBP cleavage reaction kinetics

An MBP cleavage assay was performed by mixing all components at experimental concentrations ([MBP-FUS-GFP] = 1.49 µM, [TEV protease] = 5 µM, HEPES buffer, [$MgCl_2^{total}$] = 10 mM). After addition of TEV protease and mixing with the pipette, aliquots for the different time points were separated and incubated at room temperature. The aliquots were quenched at the specified times by addition of Lämmli SDS loading buffer and DTT, followed by mixing and immediate denaturation at 95 °C for 10 min. The time span between buffer addition and placement into the pre-heated thermocycler was measured as roughly 20 seconds. After denaturation the aliquots were stored in the fridge (4 °C) over night until analyzed via SDS PAGE. The results are shown in Supplementary Fig. 17 and confirm full cleavage of the MBP tag already after 2 min.

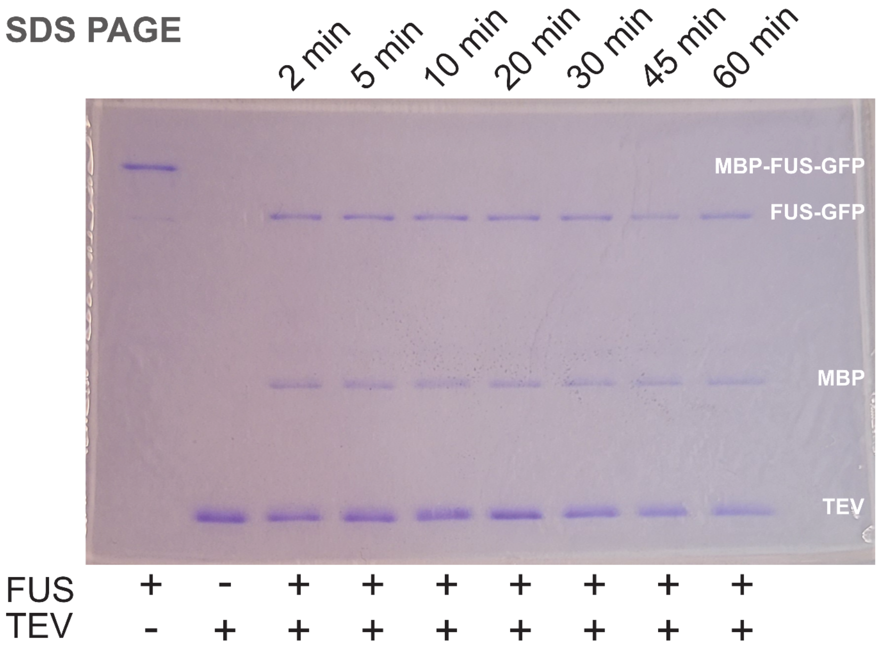

**Supplementary Fig. 17: Kinetics of the TEV-induced cleavage reaction.** SDS-PAGE: stacking gel: 4%, separation gel: 10%. Stacking voltage: 80V, 30 min. Separation voltage: 150V, 80 min. Staining: Coomassie blue. Lämmli loading buffer, Tris-Glycin running buffer.

#### 13. FUS-GFP leakage from PN

Supplementary Fig. 18 shows a representative CLSM image of $T_{20}^{Atto\,647\,N}$-PN in their final state after 2 days, as a comparison to a sample with FUS-GFP leakage as shown in Supplementary Fig. 19.

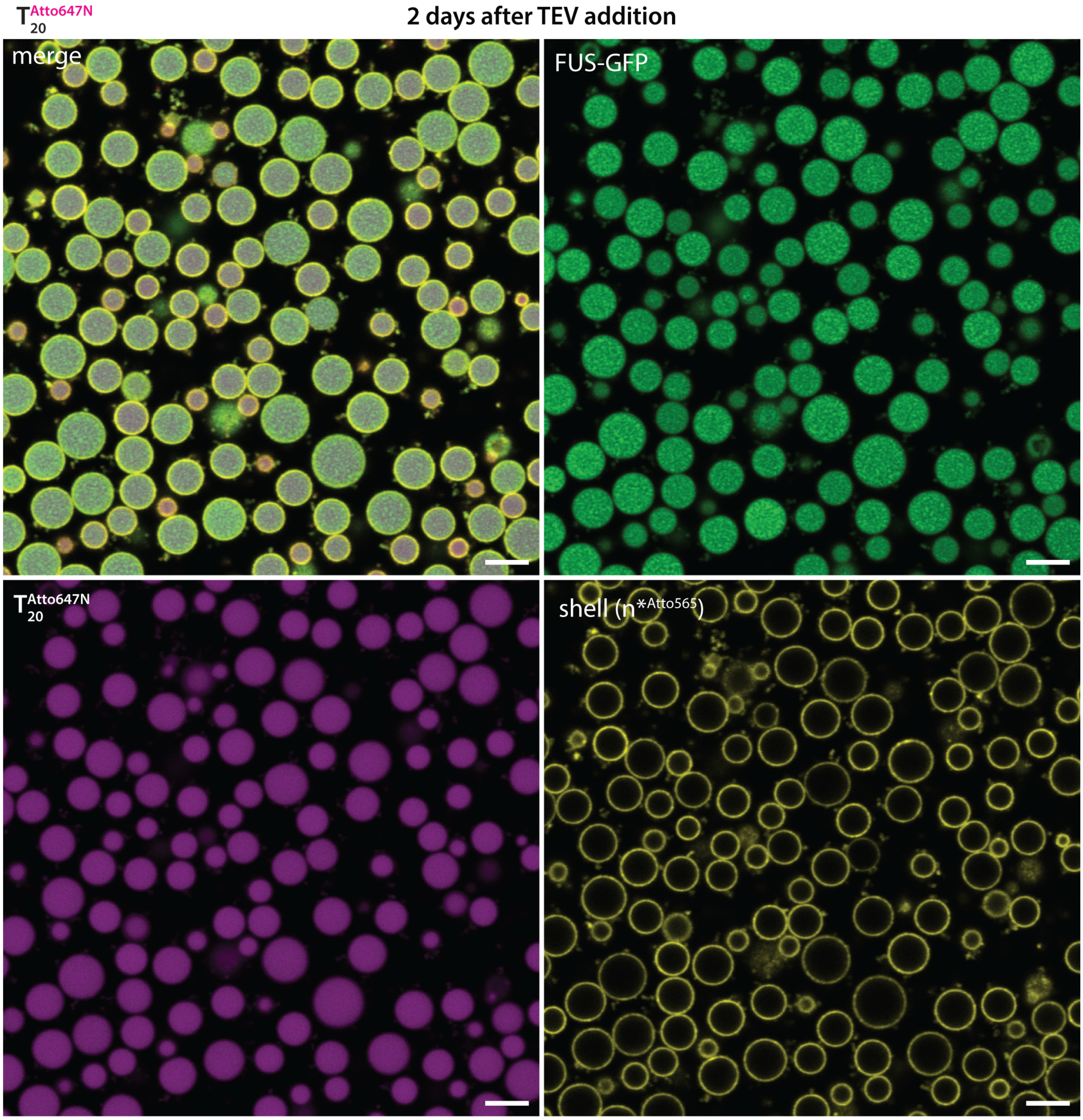

**Supplementary Fig. 18: PN remain stable for the duration of the experiment.** Confocal images of $T_{20}^{Atto\,647\,N}$-PN in their final state 2 days after TEV addition, showing minimal leakage to solution. Scale bars = 5 µm.

The overview images of m*$^{Atto647N}$-PN confirm leakage of FUS-GFP to the solution after 2 days, while the PN themselves remain stable. This poor FUS-GFP retention is not seen for the other core modifications.

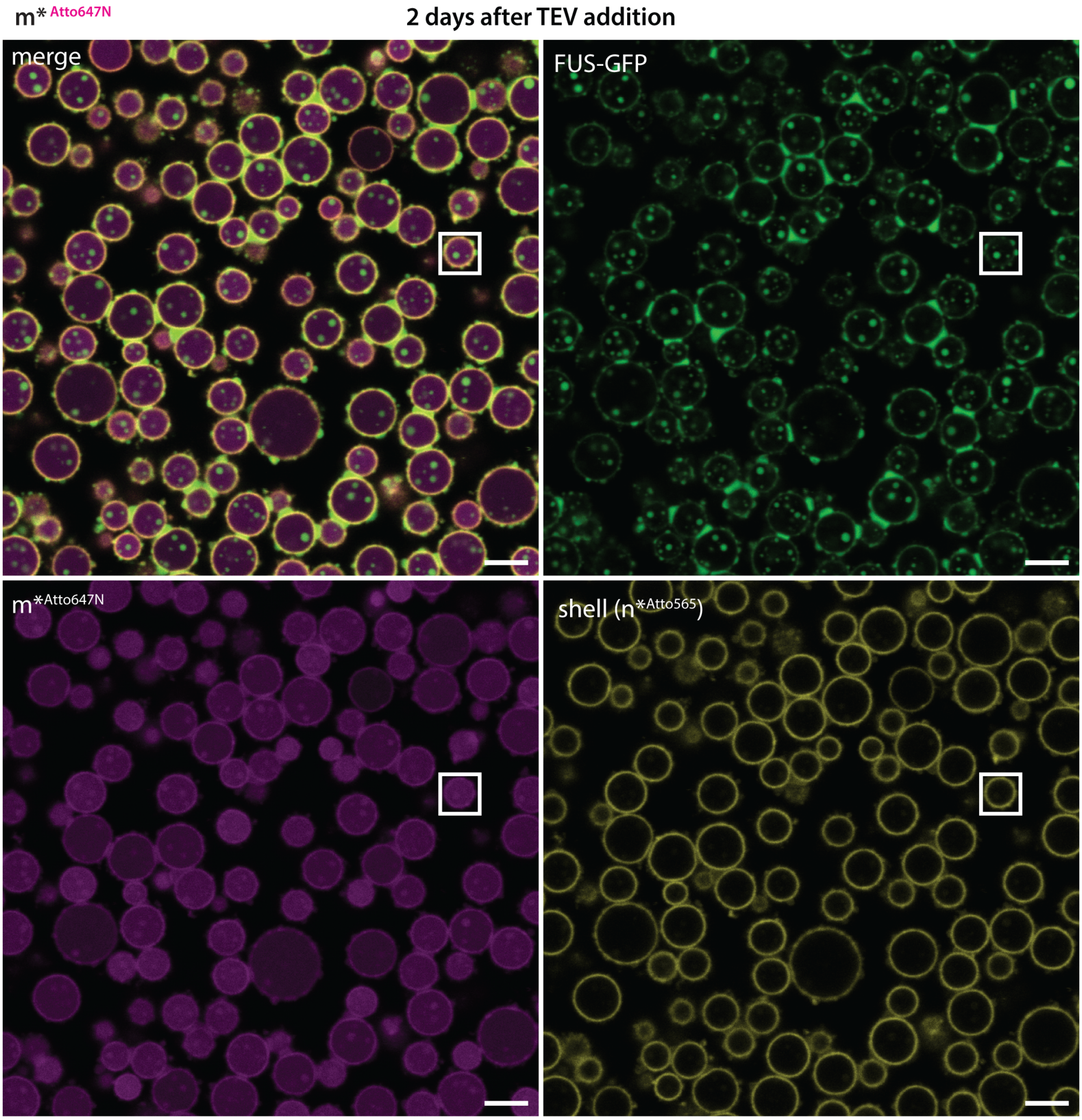

**Supplementary Fig. 19: FUS-GFP is leaking from m*Atto647N-PN over time.** The white boxes indicate the PN shown in Figure 4c. Scale bars = 5 µm.

#### 14. Covalently labelled ssDNA PN core is drawn into condensates

The following Supplementary Fig. 20 shows FUS PS within a covalently labelled PN core, to provide information about the localization of the p($A_{20}$-m) core strands for the experiment without fluorescent DNA in the core (Figure 4c, left). The labelling was performed by addition of fluorescent dNTP during RCA of the p($A_{20}$-m). Supplementary Fig. 20 confirms that the p($A_{20}$-m) core strands of pristine PN are drawn to the shell in the same way as $U_{20}$-RNA$^{Cy5}$- and $T_{20}$-m$^{Atto647N}$-PN.

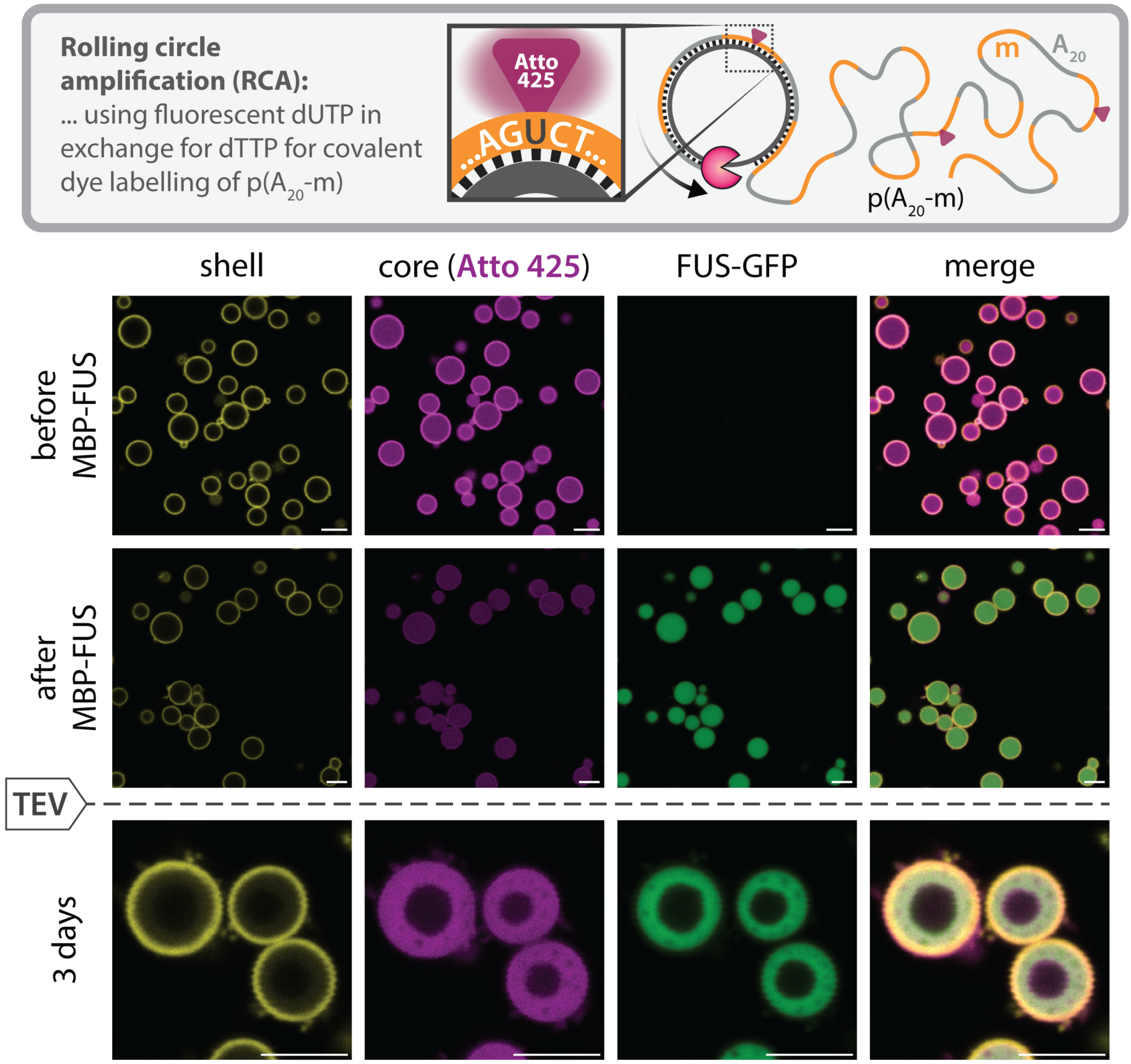

**Supplementary Fig. 20: Covalently labelled PN core without modifications.** In pristine PN, the dye-labeled pA core material is drawn into the FUS-GFP condensates forming a concentric ring at the pT interface due to $A_{20}$/$T_{20}$ affinity. Scale bars = 5 µm.

#### 15. FRAP in crosslinked PN

FRAP was conducted by bleaching the MBP-FUS-GFP channel in all samples. Recovery was only monitored for 30-45 seconds because of PN drift and unwanted bleaching during image acquisition. FRAP measurements were performed in triplicates, with different PN of the same batch. FRAP was fitted as described above and the recovery intensity after 30 s was extracted from the plots with its error.

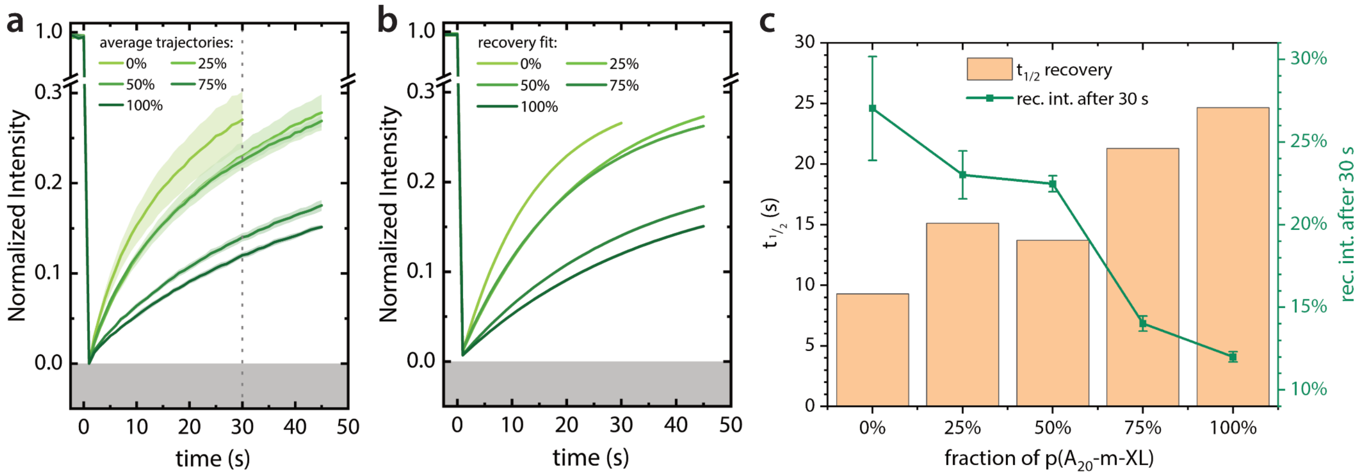

**Supplementary Fig. 21: FRAP of MBP-FUS-GFP in PN with varying amounts of crosslinker p($A_{20}$-m-XL)**. **a** average trajectories and standard deviation as a shaded region. The dotted line indicates the time point of comparison of the recovery intensity for **c**. **b** Recovery fit of the traces in **a**. **c** Comparison of $t_{1/2}$ and recovery intensity after 30 s for all samples. The error bars represent the standard deviation of the averaging step in **a**.

#### 16. Droplet assay

The established alternative to PN-experiments are so called droplet assays, where PS of the protein of interest and DNA/RNA is monitored in solution. As the name suggests, the expected outcome is droplet formation and sedimentation of the condensate phase. Typically, the assay is used to determine under which conditions, in which mixtures and with which binding partners the system forms droplets. For this reason, the result of droplet assays is often binary: does the sample show PS or not? As a comparison to our PN experiments we ran droplet assays of a strong binder (m*-RNA$^{Cy5}$) and a weak binder (m*$^{Atto647N}$) in presence of a short core sequence repeat p($A_{20}$-m)$_5$. As shown in Supplementary Fig. 22 there is hardly any morphological difference in the droplets/condensates between both binders.

Additionally, we repeated the experiment with p($A_{20}$-m)$_5$ and PEG as a crowding agent. This control experiment shows an unexpectedly strong contribution of PEG to PS, even forcing PS of the MBP-FUS-GFP protein before cleavage of the MBP tag. The cleavage reaction then triggers a second phase transition forming droplets of the dilute phase within FUS condensates, indicating that the system is situated in the inverted part of the phase diagram. Overall PEG changes the FUS phase behavior drastically, which is why it should be used with caution.

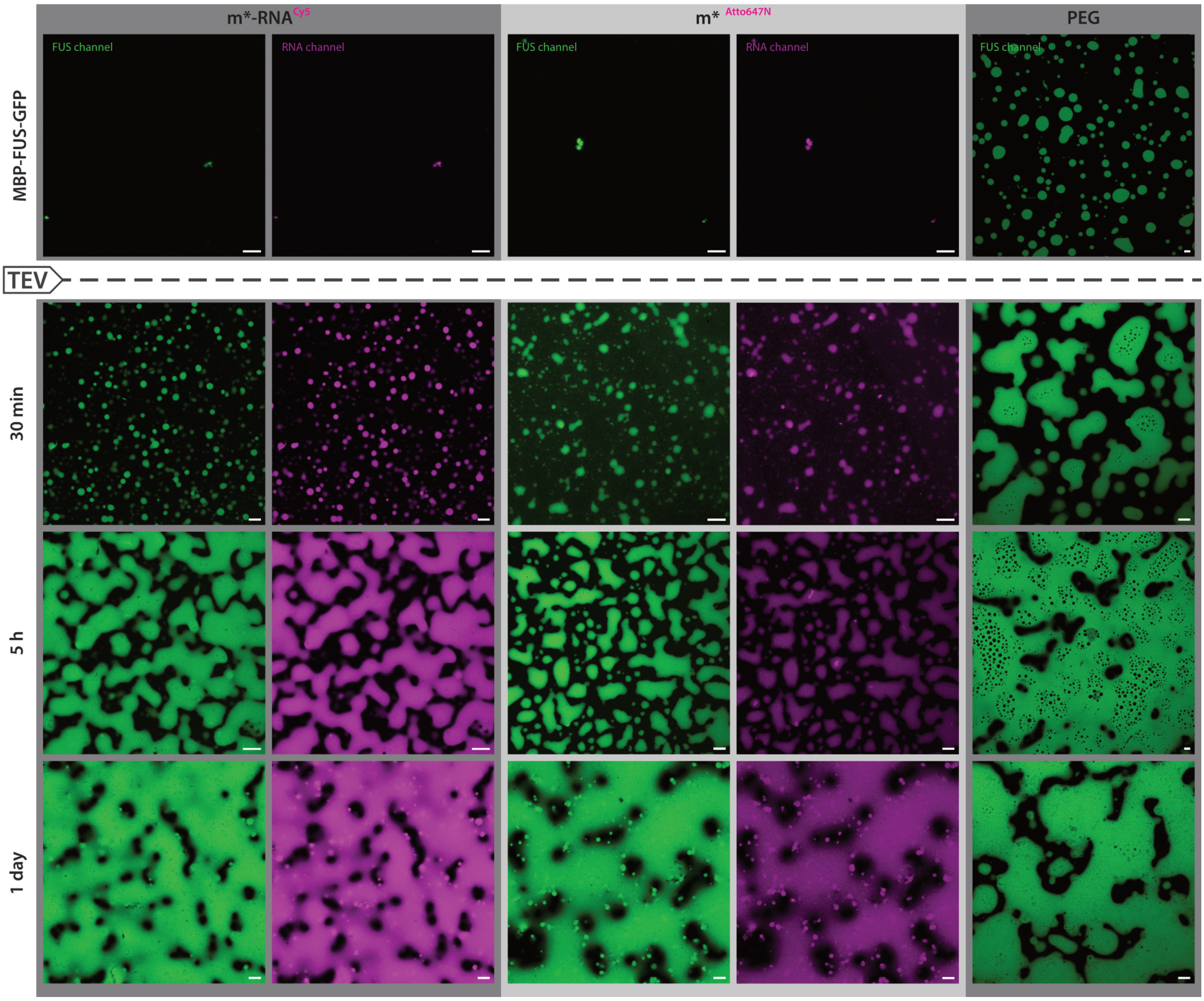

**Supplementary Fig. 22: In droplet assays FUS-GFP shows similar PS for strong (RNA) and weak (DNA) binders. Crowding agent PEG induces PS of MBP-FUS-GFP and forces FUS-GFP PS in the inverted region.** Droplet assays have been done under the same experimental conditions as PN experiments, except for the replacement of p($A_{20}$-m)$_x$ with p($A_{20}$-m)$_5$ and the omission of the shell-forming p($T_{20}$-n)$_y$. The concentration has been adapted so that the concentration of the repeating unit is the same. For the crowding experiments a 50% PEG8000 solution has been added to the final value of 10% PEG (w/v). Scale bars = 5 µm.

#### 17. Replication and statistical reporting

Unless stated otherwise, N denotes the number of individual PN compartments, droplets, or FRAP regions analysed. Time-series experiments were performed by following multiple PN in the same sample over the indicated time course. Representative images were selected from larger fields of view, which are provided in the Supplementary Information where applicable. Key qualitative trends were observed consistently across multiple PN, different PN batches, and related experimental conditions. Independent experimental repeats are explicitly stated where performed.

#### 18. Supplementary Movie legends

**Supplementary Movie 1.** Initial FUS-GFP PS in pristine PN after TEV addition. The movie shows the first 30 min of the process corresponding to Figure 1.

**Supplementary Movie 2.** Numerical simulation of PN generation from a homogeneous DNA/MBP-FUS/solvent mixture, corresponding to Figure 3a.

**Supplementary Movie 3.** Numerical simulation of DNA:FUS co-PS inside PN after MBP removal, corresponding to Figure 3b.
